# Splenic red pulp macrophages eliminate the liver-resistant *Streptococcus pneumoniae* during bloodstream infection

**DOI:** 10.1101/2024.04.26.591417

**Authors:** Yijia Huang, Zhifeng Zhao, Kunpeng Li, Xueting Huang, Xianbin Tian, Jingjing Meng, Hongyu Zhou, Jiamin Wu, Qionghai Dai, Jing-Ren Zhang, Haoran An

## Abstract

The spleen is well-known for defense against invasive infections of encapsulated bacteria, particularly *Streptococcus pneumoniae* (pneumococcus). However, the precise mechanism of the splenic anti-bacterial immunity remains elusive. Here we report that red pulp (RP) macrophages execute the splenic defense against *S. pneumoniae* in mice, with the help of natural antibodies (nAbs) and complement system. The spleen slowly but substantially cleared encapsulated pneumococci from the bloodstream in the early phase of blood infection, especially the serotypes that resist to liver filtration. Among splenic macrophage subpopulations, only the lack of RP macrophages led to the complete loss of the splenic immunity. Intravital microscopy detected direct capture and phagocytic killing of circulating pneumococci by RP macrophages in the absence of neutrophils and inflammatory monocytes. Likewise, plasma nAbs targeting pneumococcal cell wall phosphocholine and complement protein C3 were essential for RP macrophage-mediated immune clearance. Our findings have thus uncovered the long-sought immune pathway against invasive pneumococcal infections in the spleen.

**Short summary:** The spleen plays a crucial role in defending against invasive infections caused by *Streptococcus pneumoniae* (pneumococcus) and other encapsulated bacteria in humans, but the mechanisms underlying this organ-specific immunity remain largely elusive. This study has revealed that the spleen eliminates pneumococci through a coordinated interplay among red pulp macrophages, serum nAbs targeting phosphocholine and the complement system.

## INTRODUCTION

The spleen, the largest immune organ in the body, is well-known for its defense against encapsulated bacteria ^1,2^. Asplenia (the congenital or acquired absence of the spleen) greatly increases the susceptibility to overwhelming infections by several encapsulated bacteria, with an approximately 50-fold higher risk of developing a severe septic infection and a 50-70% mortality rate ^3,4^. *Streptococcus pneumoniae* (pneumococcus) is the most common causal organism associated with 50-90% of post-splenectomy infection cases, with serotype b *Haemophilus influenzae* and *Neisseria meningitidis* as the less frequent pathogens ^5^. While these bacteria are distantly related at the evolutionary scale, they are all covered by variable serotypes of polysaccharide capsules. Moreover, phosphocholine (PC) derivatives are found in the cell wall teichoic acid or C polysaccharide of *S. pneumoniae* ^6^, lipopolysaccharide of *H. influenzae* ^7^, and pilus of *N. meningitidis* ^8^. It is completely unknown why the spleen is particularly important for host resistance to these bacteria.

The spleen is the largest lymphoid organ with the close vascular connections with the digestive system through the hepatic portal system, which is formed by the spleen vein and the gastrointestinal veins ^9^. This anatomic placement is highly relevant to the two main functions of the organ: filtering senescent/damaged red blood cells and blood-borne microorganisms, and mounting innate and adaptive immune responses against pathogens ^10^. The spleen is a highly compartmentalized organ with three functionally inter-related compartments: red pulp (RP), white pulp (WP) and marginal zone (MZ) ^9^. The sponge-like RP is filled with slow-flowing blood from sinuses and cords, which is important for blood filtering by RP macrophages. The WP contains B and T lymphocytes for the maturation of B cells and antibody production. The MZ is located at the extreme periphery of the WP, and contains natural antibody-producing B cells, MZ macrophages, and metallophilic (MP) macrophages. Besides, neutrophils and monocytes are abundantly present in the spleen ^11,12^.

Despite the well-recognized importance of the spleen in host defense against circulating bacteria, the precise mechanisms behind its immune actions remain largely speculative ^5^. RP macrophages have been shown to capture blood-borne pneumococci without the capacity of phagocytic killing in mice; instead, they rely on neutrophils to kill the immobilized bacteria ^11^. In a somewhat opposite manner, a separate study has reported that RP macrophages are predominantly responsible for splenic killing of *S. pneumoniae*; neutrophils and dendritic cells are dispensable ^13^. MZ macrophages have been described to bind to pneumococcal capsules through the lectin receptor SIGN-R1 ^14,15^, which appears to contribute to host immunity to pneumococcal disease ^16,17^. Interestingly, MP macrophages seem to play an opposite role by serving as an intracellular replication site for pneumococci ^18^. Natural antibodies (nAbs) produced by marginal zone B cells have also been implicated in the opsonophagocytosis of *S. pneumoniae, H. influenzae* and *N. meningitidis* ^10^, but it is completely unclear which phagocytes in the spleen carry out the nAb-mediated phagocytosis.

The liver is also involved in the removal of invading bacteria and other foreign particles from the blood circulation beyond the spleen. Intravenously administrated vital stains and bacteria are concentrated to the spleen and liver of mammals ^19–22^, which has led to the loosely-defined concept of “reticuloendothelial system” ^23,24^. The liver is considered as the vascular “firewall” against invading bacteria in the bloodstream ^25,26^. The liver resident macrophage Kupffer cells (KCs) and sinusoidal endothelial cells are the major cell types responsible for the immune/scavenging function ^27^. Our recent works have shown that the liver is the major target organ of bacterial capsules to promote pathogen survival in the blood circulation ^28,29^. Acapsular bacteria are rapidly captured by KCs in the liver sinusoids, whereas encapsulated bacteria circumvent KC recognition and grow in the bloodstream. Furthermore, the potency of capsule-mediated immune evasion depends on the chemical structures or serotypes of capsular polysaccharides. Certain high-virulence (HV) capsular serotypes completely escape KC capture and lead to severe septic diseases, whereas other low-virulence (LV) serotypes are partially trapped in the liver and cannot cause septic disease unless they are present in the bloodstream at relatively higher concentrations. We will refer to the HV and LV serotypes as the liver-resistant and -susceptible bacteria, respectively, hereafter.

The current literature has demonstrated the importance of both the spleen and liver in host defense against bacterial infections, but it remains largely unknown how the spleen and liver divide the labor in the clearance of invading bacteria, mainly due to the lack of appropriate tools in dissecting the functional redundancy. This study takes advantage of the HV (liver-resistant) and LV (liver-susceptible) capsular types to define the specific contribution of the spleen to the clearance of encapsulated bacteria based on our previous studies ^28,29^. We found that the spleen clears the liver-resistant HV pneumococci via the orchestrated actions of RP macrophages, nAbs, and the complement system.

## RESULTS

### Spleen eliminates the liver-resistant high-virulence *S. pneumoniae*

Our previous studies have shown that the liver-resistant HV serotypes of pneumococci circumvent phagocytic capture and clearance of KCs in the liver vasculatures, leading to overwhelming septicemia and death ^28^. During that study, we noticed slow but substantial elimination of the liver-resistant pneumococci from the bloodstream of mice in the first 12 hr post intravenous (i.v.) infection, so called “eclipse” phase ^30^, although bacteremia relapsed to a lethal stage at the later phase of infection (**Fig. 1A**, *Spn*6A). In sharp contrast, liver-susceptible LV serotypes were rapidly eradicated (**Fig. 1A**, *Spn*14). To determine the extent of this anti-HV pneumococcal immunity, we performed similar infection with lower doses of *Spn*6A (10^3^-10^5^ CFU), and monitored the bacteremia kinetics (**Fig. 1B**) and survival (**Fig. 1C**). Although all the mice infected with four doses showed significant clearance of blood-borne bacteria at 12 hr, those receiving relatively higher doses displayed higher levels of bacteremia. There were undetectable blood bacteria in the group of 10^3^ CFU; mice infected with 10^4^-10^6^ CFU exhibited increasing levels of bacteremia that were proportional to the inoculum doses (**Fig. 1B**). While infection with 10^4^ CFU led to a mild bacteremia of 1,300 CFU at 12 hr, relatively higher levels of bacteremia were observed in groups of 10^5^ CFU (4,500 CFU/ml) and 10^6^ CFU (4,700 CFU/ml). Consistently, there were undetectable bacteria in mice infected with 10^3^ and 10^4^ CFU at 48 hr, substantial levels of bacteria were present in the blood circulation of mice infected with 10^5^ (3,500 CFU/ml) and 10^6^ CFU (83,000 CFU/ml). This dose-dependent bacterial clearance indicated that this uncharacterized immunity is only able to clear relatively lower levels of blood-borne HV pneumococci. This conclusion is supported by a challenge dose-dependent survival of *Spn*6A-infected mice. All the mice in the group of 10^3^ CFU survived the infection, whereas those receiving higher doses showed partial (10^4^ and 10^5^ CFU) or no (10^6^ CFU) survival (**Fig. 1C**). This trial revealed a LD_50_ of 1.3 × 10^5^ CFU for *Spn*6A in the i.v. infection route.

**Figure 1.**
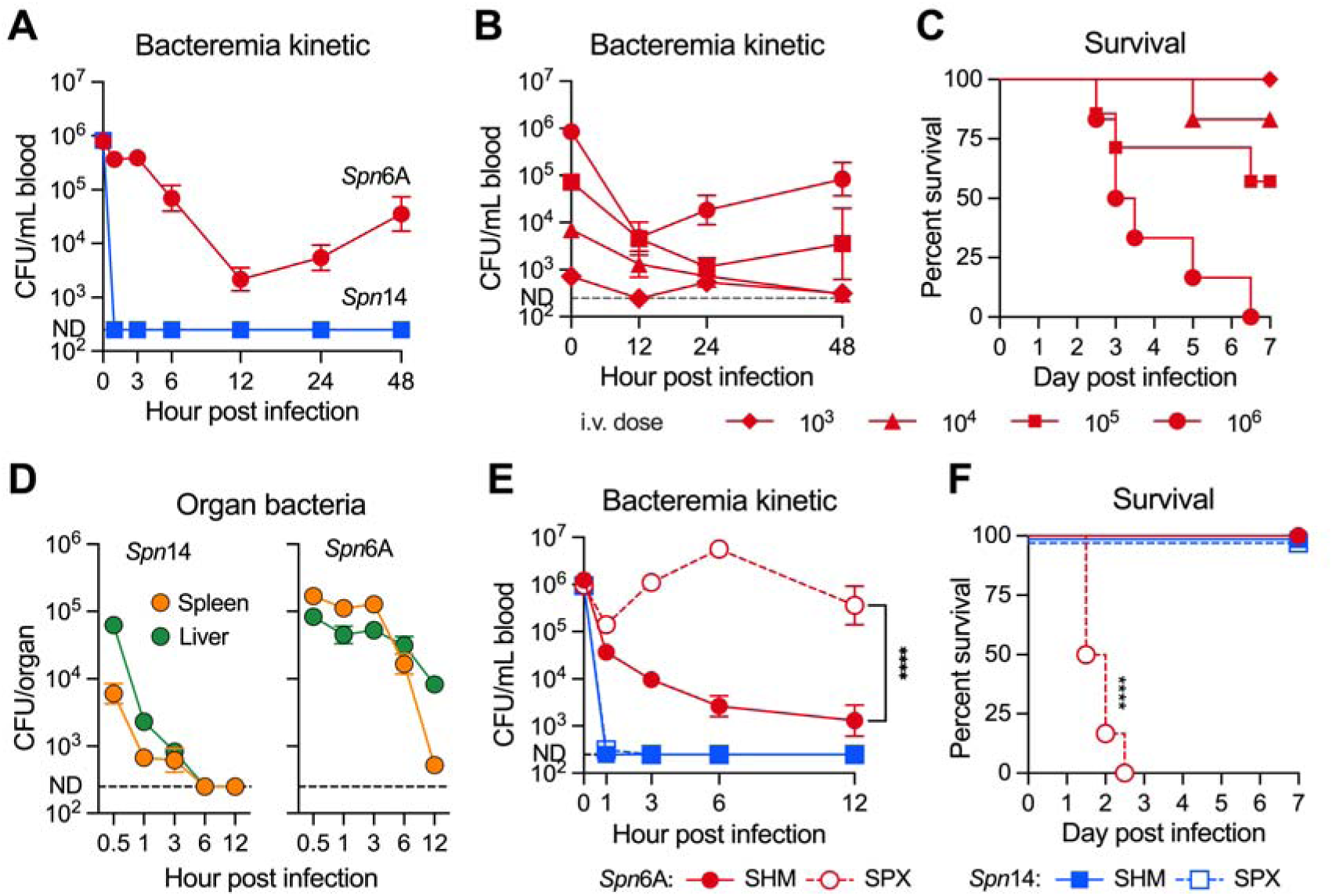
Essential function of the spleen in clearing liver-resistant pneumococci. (A) Dramatic difference in the clearance kinetics of LV and HV pneumococci from the bloodstream of mice. Blood bacteria were monitored by retroorbital plexus bleeding and CFU counting at various time points post intravenous (i.v.) infection with 10^6^ CFU LV serotype 14 (*Spn*14) or HV serotype 6A (*Spn*6A). n = 6. (B) Dose-dependent clearance of HV pneumococci. Blood bacteria were monitored in mice i.v. infected with 10^3^, 10^4^, 10^5^ or 10^6^ CFU *Spn*6A. n = 6. (C) Survival rate of mice post i.v. infection with various doses of *Spn*6A. (D) Differential distribution of LV and HV pneumococci in the liver and spleen. Bacterial loads were counted at various time points post i.v. infection with 10^6^ CFU *Spn*14 or *Spn*6A. n = 3-6 at each time point. (E) Impact of splenic removal on the clearance of HV pneumococci from the bloodstream. Blood bacteria of sham-operated (SHM) and splenectomized (SPX) mice were monitored during the first 12 hr post i.v. infection with 10^6^ CFU *Spn*14 or *Spn*6A. n = 6. (F) Survival rate of SHM and SPX mice post i.v. infection with 10^8^ CFU *Spn*14 or 10^3^ CFU *Spn*6A. n = 6. All data were pooled from two independent experiments. Dotted line indicates the detection limit (A, B, D, and E). Significance was compared by two-way ANOVA (E) and log-rank test (F). **** *P* < 0.0001.

To determine how the liver-resistant pneumococci are cleared, we compared the levels of LV and HV pneumococci in the major organs at various time points post i.v. inoculation. As reported in our previous work ^28^, *Spn*14 bacteria were predominantly trapped in the liver at the onset of blood infection and became undetectable at 6-12 hr, but *Spn*6A pneumococci were more abundant in the spleen than liver in the first 3 hr and reduced to barely detectable level at 12 hr (**Fig. 1D**). Consistently, mice lacking KCs did not show obvious deficiency in clearing *Spn*6A in the first 12 hr of blood infection (**Fig. S1**). These results indicated that HV pneumococci are eliminated by a liver-independent immune mechanism in the spleen. To test this hypothesis, we compared bacteremia kinetics between asplenic and normal mice. In contrast to dramatic reduction in *Spn*6A bacteremia in sham-operated (SHM) mice in the first 12 hr of blood infection, splenectomized (SPX) mice were no longer able to control the HV bacteria, with sustained bacteremia during this period (**Fig. 1E**). The importance of the spleen in clearing HV pneumococci was also evidenced by the hyper-susceptibility of SPX mice to *Spn*6A; all SPX mice died in 3 days post i.v. infection with an otherwise non-lethal dose (10^3^ CFU) (**Fig. 1F**). Consistent with liver-based immunity against LV bacteria ^28^, SPX mice were full competent in clearing *Spn*14 (**Fig. 1E**), and withstood i.v. infection with a high dose (10^8^ CFU) of *Spn*14 (**Fig. 1F**). These results demonstrated that the spleen is the major immune organ to eliminate the liver-resistant HV pneumococci in the early phase of blood infection.

### RP macrophages are essential for splenic elimination of the liver-resistant pneumococci

To identify the splenic immune cells that are responsible for clearing HV pneumococcus, we first tested the splenic macrophages by selective depletion with clodronate liposomes ^31^, because opposing roles of splenic macrophages have been reported in terms of their roles in anti-bacterial immunity ^11,13,18^. As compared with control mice, depleting both MZ and MP macrophages with a low-dose of clodronate liposomes (LDL) did not yield significant impact on the clearance of *Spn*6A in the first 12 hr post i.v. infection (**Fig. 2A**), suggesting that MZ and MP macrophages are dispensable for splenic clearance of HV pneumococci. In contrast, additional depletion of RP macrophages with a higher dose of clodronate liposomes (HDL) resulted in dramatic impairment in *Spn*6A clearance (**Fig. 2A**). Since the extent of immune dysfunction in HDL-treated mice was highly similar to what was observed in SPX mice (**Fig. 1E**), this result strongly suggested that RP macrophages are the major immune cells for clearing the liver-resistant pneumococci.

**Figure 2.**
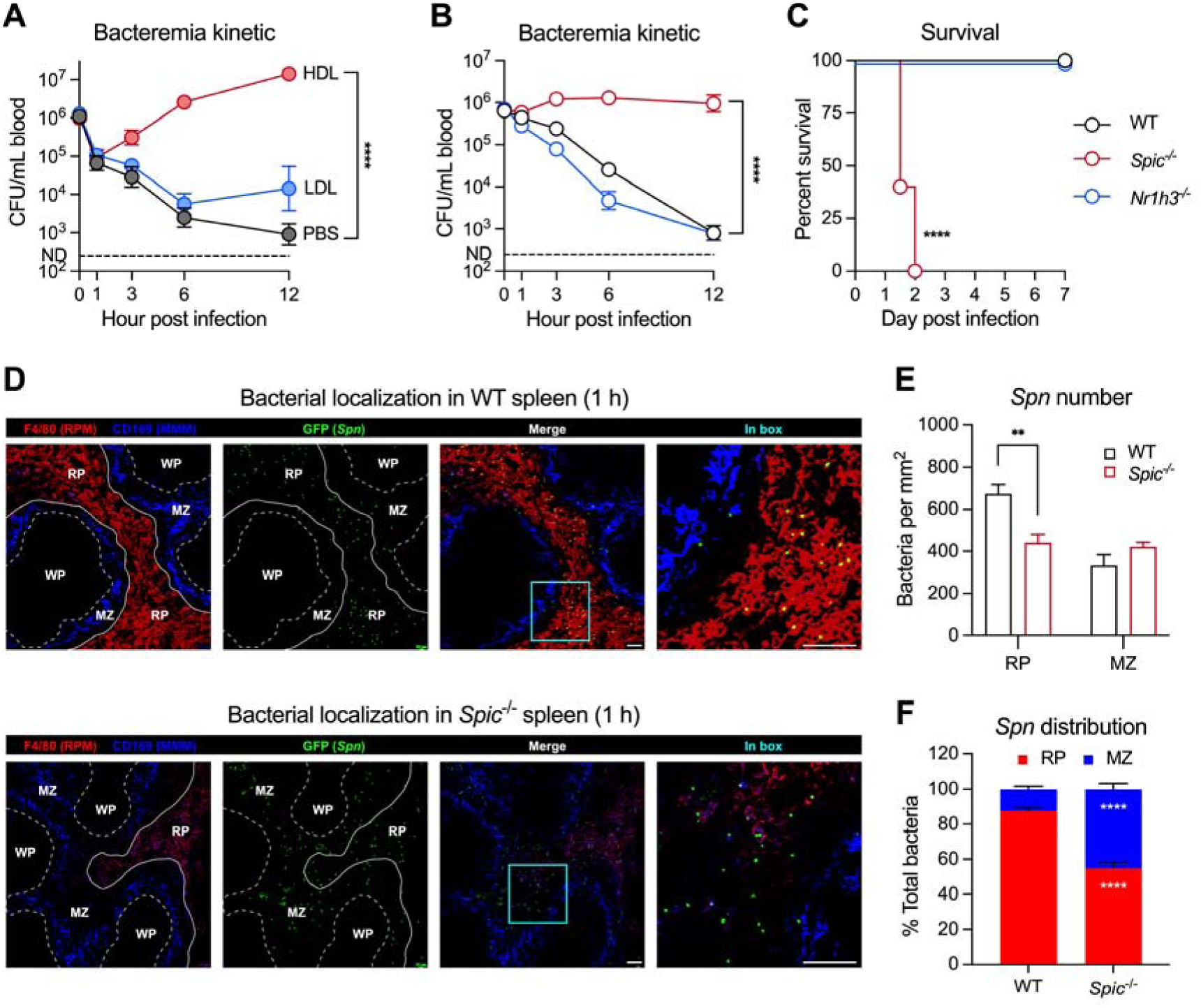
The importance of RP macrophages in splenic control of HV pneumococci. **(A)** Impact of macrophage depletion on the clearance of HV pneumococci. Blood bacteria were monitored in mice that were pretreated with low-dose clodronate liposome (LDL) or high-dose clodronate liposome (HDL) and i.v. infected with 10^6^ CFU *Spn*6A. n = 3-6. **(B)** Essentiality of RP macrophages for the removal of HV pneumococci. Bacterial kinetics in the bloodstream were monitored in RP macrophage-deficient *Spic*^-/-^ mice or *Nr1h3*^-/-^ mice lacking both MZ and MP macrophages post i.v. infection with 10^6^ CFU *Spn*6A. n = 6. **(C)** Survival of *Spic*^-/-^ and *Nr1h3*^-/-^ mice post i.v. infection with 10^3^ CFU *Spn*6A. n = 6. **(D)** Representative immunofluorescent images to show bacterial trapping in splenic RP of WT and *Spic*^-/-^ mice. Mice were i.v. infected with 10^7^ CFU *Spn*6A-GFP (green) and splenic sections were prepared at 1 hpi. RP and MP macrophages were stained by AF647 anti-F4/80 (red) and AF594 anti-CD169 (blue), respectively. **(E and F)** Quantitative analysis of the imaging data in (D). Bacteria in 1 mm^2^ per area of the RP and MZ were counted (E). Normalized bacteria distribution (F) was calculated by multiplying bacterial number per area by the total area of RP or MZ in the splenic section. n = 5 random fields of view (FOVs). Scale bar, 40 μm. Data were pooled (A to C) or representative results (D to F) from two independent experiments. Dotted line indicates the detection limit (A and B). Significance was compared by two-way ANOVA (A, B, E, and F) and log-rank test (C). ** *P* < 0.01, **** *P* < 0.0001.

We verified the unique importance of RP macrophages in splenic clearance of HV pneumococci using *Spic*^-/-^ mice (with a deficiency in RP macrophage development) ^32^ or *Nr1h3*^-/-^ mice lacking both MZ and MP macrophages ^33^. *Spic*^-/-^ mice showed a similar extent of severe functional deficiency as HDL-treated and SPX mice in clearing HV pneumococci, with sustained bacteremia in the first 12 hr post i.v. inoculation of *Spn*6A, whereas *Nr1h3*^-/-^ mice displayed a virtually normal pattern of bacterial clearance as wildtype (WT) control (**Fig. 2B**). Loss of RP macrophages also led to uncontrolled bacterial replication in the spleen and liver (**Fig. S2A**). In a similar fashion, all of *Spic*^-/-^ mice succumbed to i.v. infection with non-lethal dose of *Spn*6A (10^3^ CFU) in 2 days, but there was no mortality in *Nr1h3*^-/-^ mice (**Fig. 2C**). These results demonstrated that RP macrophages are the major splenic macrophages for eliminating HV pneumococci in the “eclipse phase” of blood infection.

We further characterized pneumococcus-macrophage interaction by analyzing the spleen sections with immunofluorescence microscopy. Consistent with the dominant role of RP macrophages in bacterial clearance, the GFP-labeled *Spn*6A were predominantly co-localized with F4/80-labeled RP macrophages in the RP of WT mice (**Fig. 2D**, upper panel). Relatively fewer bacteria were found in the MZ where CD169-labeled MP macrophages were located. Pneumococcal dominance in RP over MZ was diminished in *Spic*^-/-^ mice (**Fig. 2D**, lower panel), in terms of bacterial numbers per area of anatomical structure (**Fig. 2E**). After normalized by the area size of each compartment in the spleen section (**Fig. S2B**), approximately 90% bacteria were constrained in RP in WT mice, while the bacteria were uniformly distributed in the RP and MZ in *Spic*^-/-^ mice (**Fig. 2F**). These microscopic observations provided additional evidence to support the unique anti-pneumococcal activity of RP macrophages in the spleen.

### RP macrophages are capable of phagocytic killing of pneumococci

To understand how RP macrophages eradicate HV pneumococci, we carried out systemic analyses of the anti-bacterial function of splenic phagocytes. We first determined the roles of neutrophils and inflammatory monocytes (IM), given the previous report that neutrophils are required to kill pneumococci captured by RP macrophages ^11^. Flow cytometry analysis showed dramatic increase in neutrophils and IMs in the first 6 hr of HV infection (**Fig. 3A and S3A**), which was effectively diminished by antibody depletion (**Fig. S3B**). In contrast to dramatic impairment of RP macrophage-deficient mice in bacterial clearance, depleting neutrophils with 1A8 antibody did not yield obvious impact on bacteremia kinetics post i.v. inoculation (**Fig. 3B**). Likewise, *Ccr2*^-/-^ mice with the deficiency in IM recruitment ^34^ did not exhibit significant defect in pneumococcal clearance (**Fig. 3B**). Surprisingly, simultaneous depletion of both neutrophils and IMs using Gr1 antibody even enhanced pneumococcal clearance at the early time points (3 and 6 hrs). Although Gr1-treated mice displayed functional deficit at the later stage (12 hr), the blood bacteria were still 15-fold lower than that in *Spic*^-/-^ mice (**Fig. 3B**), as well as 150- and 35-fold lower bacterial burdens in the spleen and liver (**Fig. S3C and S3D**). This result showed that neutrophils and IMs play a minor role in splenic elimination of HV pneumococci at the early stage of blood infection.

**Figure 3.**
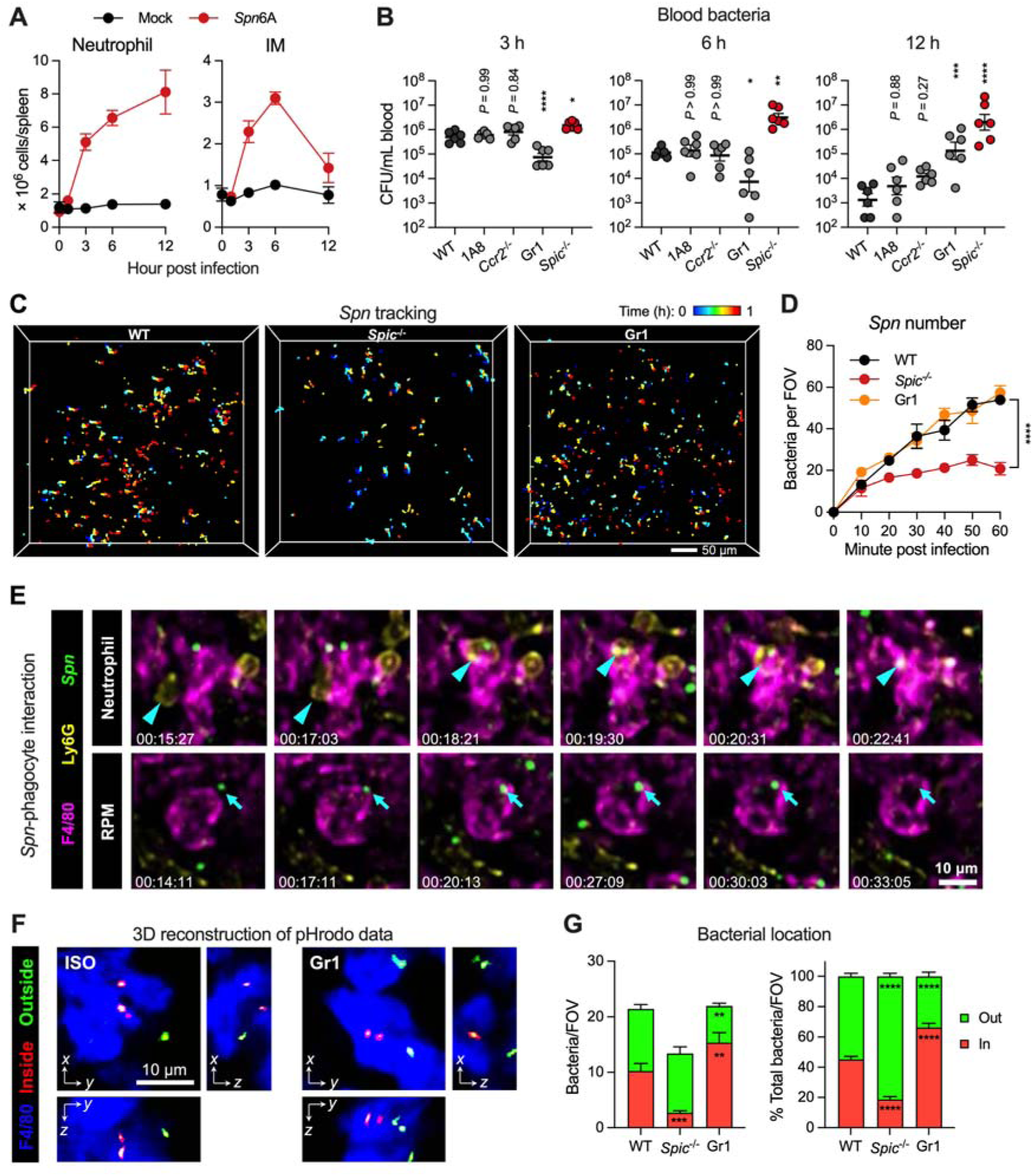
Neutrophil-independent elimination of HV pneumococci by RP macrophages. **(A)** Infiltration of neutrophils and IMs in the spleen during pneumococcal infection. The kinetics of absolute numbers of neutrophils and IMs were assessed by flow cytometry post i.v. infection with 10^6^ CFU of *Spn*6A. Mock infected mice were i.v. injected with 100 μl of Ringer’s solution. n = 3. **(B)** Impact of circulating phagocyte depletion on the clearance of HV pneumococci. Bacterial loads in the blood were counted at 3, 6, and 12 hpi with 10^6^ CFU of *Spn*6A and compared among phagocyte-deficient mice. 1A8 or Gr1 antibodies depleted neutrophils or both neutrophils and IMs, respectively; *Ccr2*^-/-^ mice were deficient in IM infiltration; *Spic*^-/-^ mice lacked RP macrophage; ISO, isotype control. n = 6. **(C)** 2pSAM illustration of pneumococcal tracking in the spleens of WT, *Spic*^-/-^, and Gr1 mice post i.v. infection with 5 × 10^6^ CFU of *Spn*6A-GFP (Movie S2). Pneumococci trapped in the field for at least 1 minute were recorded during the first hour post infection. Scale bar, 50 μm. **(D)** Number of trapped pneumococci in the field during the first hour as monitored by 2pSAM in (C). n = 4 random FOVs. **(E)** Representative consecutive 2pSAM imaging to show the neutrophil-dependent elimination (upper panel, Movie S3A) and direct elimination by single RP macrophage (lower panel, Movie S3B) of *Spn*6A in the spleen of WT mice. Arrows and arrow heads indicate *Spn* cells and neutrophils, respectively. Scale bar, 10 μm. **(F)** Representative 3D reconstruction of 2pSAM images to illustrate the uptake of pneumococci by RP macrophages. Internalization of *Spn*6A cells were labelled with pHrodo Red and i.v. injected into ISO and Gr1-treated mice. Internalization of bacteria was indicated by the activation of pHrodo Red under acid environment, i.e. in the phago-lysosomes of RP macrophages. Images were obtained at 1 hpi. Scale bar, 10 μm. **(G)** Quantification of pneumococcal uptake by RP macrophages. The number (left) and ratio (right) of intra- and extracellular pneumococci in the spleens of WT, *Spic*^-/-^, and Gr1-treated mice were analyzed at 1 hpi. n = 5-10 random FOVs. Data were pooled (B) or representative results (A, C to G) from two independent experiments. Significance was compared by one-way (B) or two-way ANOVA (D and G) test. * *P* < 0.05, ** *P* < 0.01, *** *P* < 0.001, **** *P* < 0.0001.

To understand the process of bactericidal action of RP macrophages, we applied a two-photon synthetic aperture microscopy (2pSAM) system to visualize the *in situ* pathogen-macrophage interaction during blood infection. The newly developed 2pSAM enables long-term imaging of deep tissues at a millisecond scale while maintains 1,000-fold reduction in phototoxicity as compared to traditional two-photon microscopy ^35^. We first observed the behaviors of RP macrophages and neutrophils in response to pneumococci in the spleen of normal mice. In contrast to fast and unidirectional movement of HV encapsulated bacteria in the liver sinusoids ^28,29^, pneumococci were flowing in the splenic RP without clear direction (**Fig. 3C and 3D; Movie S1**), which agrees with the open blood system in the RP. Likewise, bacterial capture by RP macrophages occurred in a much less dramatic manner as compared with the swift and firm bacterial capture by KCs in the liver sinusoids. Many bacteria initially touched the immune cells and then moved away in a stop-go cycle (**Fig. S3E**). In line with the flow cytometry analysis, we observed an evident infiltration of neutrophils within 2 hpi (**Fig. S3F; Movie S1**). Remarkably, the development of RP macrophages is impaired as indicated by a shrink of F4/80-positive area in *Spic*^-/-^ spleen, in which the RP showed a significantly lower level of immobilized pneumococci (**Fig. 3C and 3D; Movie S2**). However, the gradual capture of *Spn*6A was competent in the spleens of neutrophil- and IM-depleted mice (**Fig. 3C and 3D; Movie S2**). This real-time observation further demonstrated the pivotal role of RP macrophages in eliminating the liver-resistant bacteria.

Next, we followed the destiny of the bacteria once being captured by RP macrophages. As previously reported ^11^, neutrophils were abundantly present in the RP and occasionally plucked the pneumococci from RP macrophage surface (**Fig. 3E upper; Movie S3A**). However, the vast majority of RP macrophages engulfed the immobilized pneumococci at the site far away from migrating neutrophils. The GFP signals were finally immersed in F4/80-positive cells, indicating intracellular digestion of the bacteria by RP macrophages (**Fig. 3E lower; Movie S3B**). Phagosome maturation after fusion with lysosome is the major way for intracellular killing of ingested bacteria, which accompanied with acidification and oxidation in phago-lysosome ^36^. We thus tested whether the disappeared pneumococci were killed in the phago-lysosome of RP macrophages by using the indicative dye pHrodo Red ^37^. As expected, the pneumococcus-loaded probes were activated as early as 30 min post inoculation in both normal and Gr1-treated mice (**Fig. 3F and S3G**). Notably, more pneumococci were embedded in the phago-lysosome of RP macrophages when neutrophils and IMs were depleted (66% inside) compared to the control (45% inside) (**Fig. 3G**), which was consistent with the accelerated bacterial clearance from the blood (3 and 6 hrs, **Fig. 3B**). However, only 18% pneumococci were internalized when RP macrophages were defective (**Fig. 3G**). Consistent with the marginal impact of neutrophil depletion on splenic clearance of HV pneumococci, these imaging data demonstrated that RP macrophages are capable of phagocytic killing of capture pneumococci without the assistance of neutrophils.

### RP macrophages utilize natural antibodies (nAbs) to clear the liver-resistant pneumococci

Our recent work has revealed that the liver KCs capture LV pneumococci by recognizing bacterial capsules through specific receptors ^28^. We thus reasoned that RP macrophages might employ a similar strategy to capture HV pneumococci. To test this hypothesis, we conducted affinity screening by incubating beads coated with capsular polysaccharides of HV pneumococcal serotypes with enriched membrane proteins of murine RP macrophages. However, our repeated trails did not consistently identify any RP macrophage membrane proteins potentially interacting with the HV capsules. This result implied that RP macrophages use an indirect manner to capture HV pneumococci, i.e. with the help of opsonins like antibody and complement proteins.

Given the fact that HV serotypes are highly proliferative in the blood of mice with genetic deficiency in natural antibody production ^38,39^, we tested if nAbs are involved in RP macrophage-mediated pneumococcal clearance using *µMT* mice, which lack mature B cells and antibodies ^40^. *µMT* mice showed severely compromised clearance of *Spn*6A, with stably sustained bacteremia in the first 6 hpi (**Fig. 4A**), although the functional deficit in *µMT* mice was less pronounced than in asplenic mice (**Fig. 1E**) and *Spic*^-/-^ mice (**Fig. 2B**) during 6-12 hpi. We next assessed if purified serum antibodies enhance pneumococcal clearance in *µMT* mice. ELISA test revealed that normal murine serum contains substantial level of IgM antibodies and, to a less extent, of IgG antibodies targeting *Spn*6A pneumococci (**Fig. S4A**). Pre-opsonization with purified natural IgM (nIgM) or nIgG restored the capacity of *µMT* mice to clear *Spn*6A in a dose-dependent manner (**Fig. 4B**). Moreover, the nIgM antibodies were more efficient than nIgG in promoting pneumococcal clearance especially in the early time points, e.g. 3 and 6 hpi (**Fig. 4B**). These results suggested that RP macrophages utilize serum antibodies to clear the liver-resistant pneumococci.

**Figure 4.**
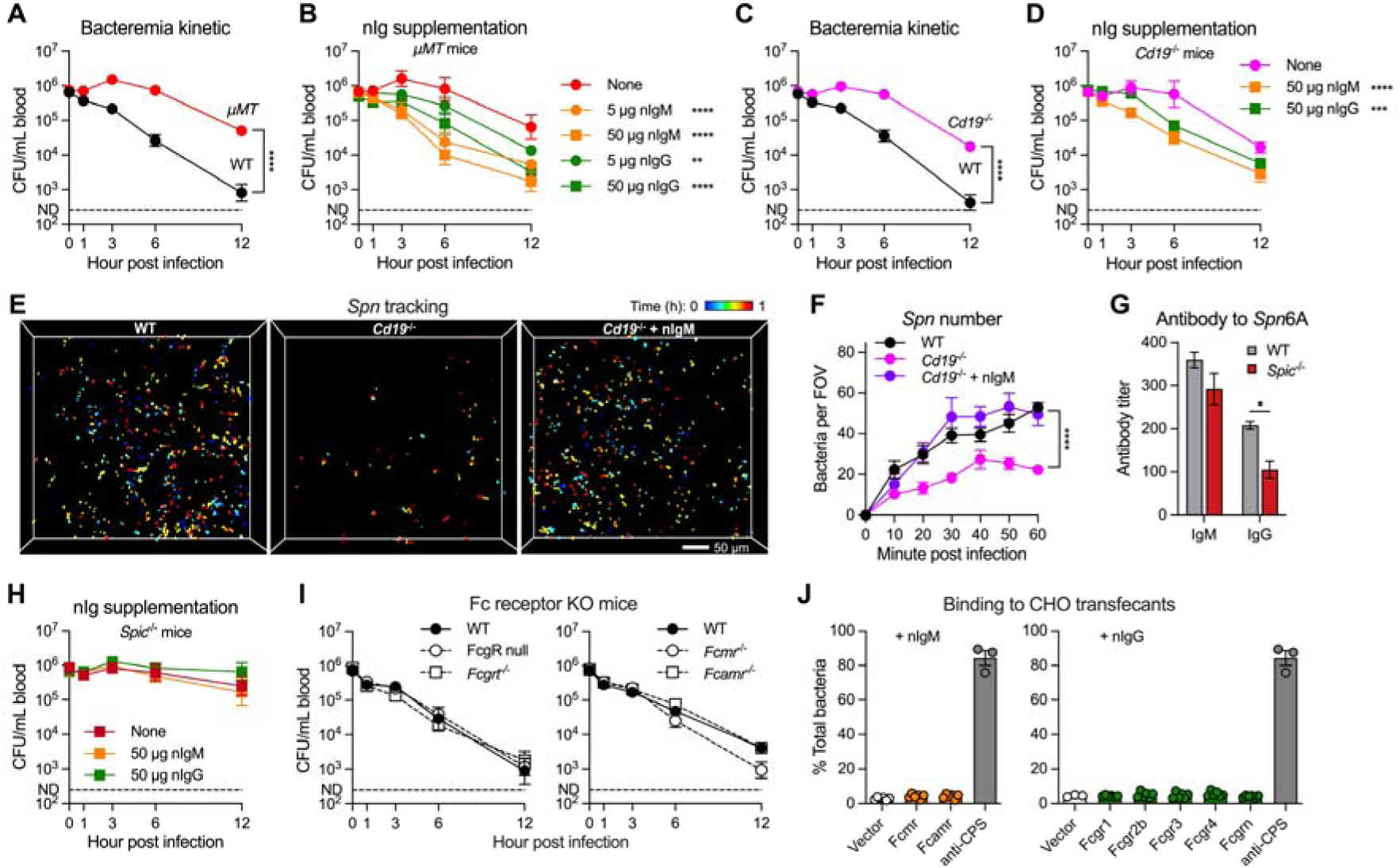
The essential role of natural antibodies in splenic clearance of HV pneumococci. **(A)** Early-phase bacteremia kinetic in WT and antibody-null (*µMT*) mice post i.v. infection with 10^6^ CFU of *Spn*6A. n = 6. **(B)** Dose-dependent promotion of *Spn*6A clearance by purified natural IgM (nIgM) and nIgG in *µMT* mice. n = 3-6. **(C)** Early-phase bacteremia kinetic in WT and B1 cell-deficient (*Cd19*^-/-^) mice post infection with 10^6^ CFU of *Spn*6A. n = 3-6. **(D)** Promotion of *Spn*6A clearance by nIgM and nIgG in *Cd19*^-/-^ mice. n = 3. **(E)** 2pSAM illustration of pneumococcal tracking in the spleens of WT and *Cd19*^-/-^ mice post infection with 5 × 10^6^ CFU of native *Spn*6A-GFP or pre-opsonized with 50 µg nIgM (*Cd19*^-/-^ + nIgM, Movie S4). Scale bar, 50 μm. **(F)** Number of trapped pneumococci in the field during the first hour as monitored by 2pSAM in (E). n = 4 random FOVs. **(G)** Titers of anti-*Spn*6A natural IgM and IgG in the serum of WT and *Spic*^-/-^ mice. n = 3. **(H)** Requirement of RP macrophages for natural antibodies in clearing HV pneumococci. Infection with nIgM- or nIgG-opsonized *Spn*6A were performed in *Spic*^-/-^ mice and bacteremia kinetics were monitored. n = 3-6. **(I)** Dispensable role of well-known Fc receptors for HV pneumococcal clearance. Bacteremia kinetics were monitored in WT and all known Fc gamma receptors-deficient (FcgR null and *Fcgrt*^-/-^) and putative Fc miu receptor-deficient (*Fcmr*^-/-^ and *Fcamr*^-/-^) mice post infection with 10^6^ CFU of *Spn*6A. n = 3-6. **(J)** *In vitro* binding of T15 IgM- or IgG3-coated HV pneumococci to CHO transfectants. *Spn*6A was added at MOI of 1 and incubated with the CHO cells for 1 hr in the presence of 50 µg/ml T15 antibodies; data are presented as the percentage of cell-associated bacteria out of the total CFU. Anti-CPS IgG1 was used as positive control for comparison. n = 3-6. Data were pooled (A to D, H, and I) or representative results (E to G, and J) from two independent experiments. Significance was compared by two-way (A, C, F, and G) or one-way (B, D, and H) ANOVA test. * *P* < 0.05, ** *P* < 0.01, *** *P* < 0.001, **** *P* < 0.0001.

To distinguish the role of nAbs produced by B1 cells from antibodies generated by antigen-stimulated B2 cells, we assessed pneumococcal clearance in B1-deficient *Cd19*^-/-^ mice, which lack nAbs ^41^. The *Cd19*^-/-^ mice showed similar functional deficiencies in producing anti-pneumococcal antibodies (**Fig. S4A**) and clearing *Spn*6A from the bloodstream as *µMT* mice (**Fig. 4C**). The bacterial clearance was substantially recovered when the *Spn*6A pneumococci were pre-coated with purified nIgM and, to a less extent, purified nIgG (**Fig. 4D**). The functional impairment of bacterial clearance was further confirmed by intravital 2pSAM. The number of trapped *Spn*6A-GFP in the spleens of *Cd19*^-/-^ mice was significantly fewer than that in WT controls within 1 hr of observation (**Fig. 4E and 4F; Movie S4A**). This immune defect was substantially reversed by opsonization of the *Spn*6A with 50 μg purified nIgM (**Fig. 4E and 4F; Movie S4B**). We finally investigated the necessity of RP macrophages for natural antibody-mediated pneumococcal clearance using *Spic*^-/-^ mice, which produce normal level of natural IgM but less IgG targeting *Spn*6A cells (**Fig. 4G**). Nevertheless, pre-coating with high doses of purified nIgM or nIgG did not improve bacterial clearance in *Spic*^-/-^ mice (**Fig. 4H**). These results showed that B1 cell-derived nAbs are essential for RP macrophage-mediated clearance of HV pneumococci in the spleen.

We next investigated how nAbs enable RP macrophages to clear circulating pneumococci. Current understanding of the antibody effector mechanisms for bacterial elimination include direct recognition by corresponding Fc receptors ^42^ and activation of the complement system via classical pathway ^43^. Our recent work has revealed that vaccine-elicited anti-capsule antibodies enable liver KCs to capture circulating pneumococci through multiple Fc receptors ^44^. Thus, we tested whether RP macrophages employ a similar mechanism to capture HV pneumococci by investigating the roles of Fc receptors. Our proteomic analysis detected the expression of FcγRIIB, FcγRIII, FcγRIV, and FcRn on murine RP macrophages, four of the five known FcγRs in mice (**Fig. S4B**). Contrary to our expectation, the FcγR null mice with complete knockout of FcγRI/IIb/III/IV ^45^, as well as FcRn KO (*Fcgrt*^-/-^) mice displayed comparable levels of pneumococcal clearance as WT counterparts (**Fig. 4I**, left panel). Similar results were observed using mice lacking FcµR or FcαµR (**Fig. 4I**, right panel), the two known IgM receptors ^46,47^. To address the potential functional redundancy among these Fc receptors, we conducted a gain-of-function approach by overexpressing the individual receptor in CHO cells and assessing pneumococcal binding to the transfectants in the presence of T15 monoclonal antibodies. However, stable expression of the two FcμRs or five FcγRs did not lead to a significant increase in adhesion (less than 5%) of T15 IgM- or IgG3-opsonized *Spn*6A to CHO cells, compared to the pronounced enhancement (85% binding) by anti-capsule antibodies (**Fig. 4J**). Collectively, the *in vivo* and *in vitro* data suggested that anti-PC nAbs contribute to HV pneumococcal clearance through activating the complement system, instead of engaging antibody receptors as the capsule-targeting antibodies ^44^.

### Protective nAbs target pneumococcal cell wall phosphocholine

To ascertain how nAbs enable RP macrophages to clear HV pneumococci, we characterized pneumococcal antigens that are recognized by the antibodies. Given our finding that the splenic anti-bacterial immunity is effective against multiple HV serotypes of *S. pneumoniae* (see below), we reasoned that protective nAbs must target a conserved antigen beyond the capsule structure. Consistently, administration of purified serotype 6A CPS before i.v. inoculation of *Spn*6A showed little impact on the bacterial clearance during the “eclipse phase” (**Fig. S5A**). These results suggested that RP macrophages recognize non-capsular ligands on HV pneumococci. The cell wall phosphocholine (PC) or C polysaccharide is the only known pneumococcal antigen that is recognized by protective nAbs ^38,48^. We verified the existence of anti-PC antibodies in normal mouse serum by ELISA, in which the IgM and IgG titers were significantly lower using PC-free *Spn*6A whole cells as antigen compared to intact pneumococci (**Fig. 5A**). The assessment also revealed substantial amounts of anti-PC IgM and IgG in normal serum (**Fig. 5A**) and purified nAbs (**Fig. S5B**).

**Figure 5.**
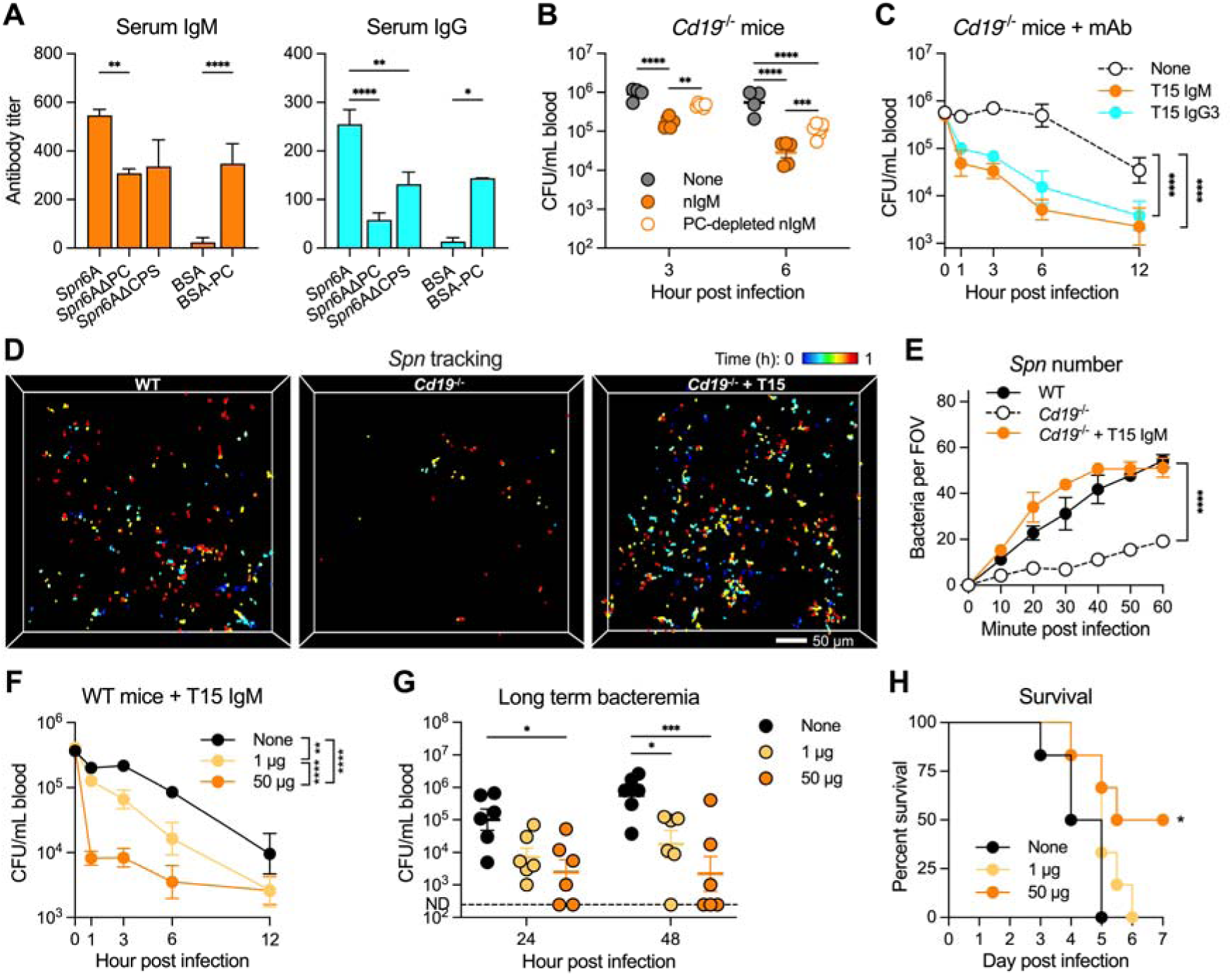
Pneumococcal cell wall PC as a target antigen of natural antibodies. **(A)** Detection of anti-PC natural antibodies in normal murine serum. Titers of IgM (left) and IgG (right) were measured by ELSIA using *Spn*6A whole cells, *Spn*6A lacking PC or CPS, and BSA-conjugated PC as antigens. n = 3. **(B)** Importance of anti-PC natural antibodies for HV pneumococcal clearance. Blood bacterial loads were counted in *Cd19*^-/-^ mice post infection with 10^6^ CFU of *Spn*6A that untreated, pre-opsonized with 50 µg native nIgM or PC resin-absorbed nIgM. n = 4-6. **(C)** Promotion of HV pneumococcal clearance by monoclonal anti-PC IgM and IgG3. Bacteremia kinetics were monitored in *Cd19*^-/-^ mice post infection with 10^6^ CFU of *Spn*6A pre-opsonized with 5 µg monoclonal T15 anti-PC IgM and IgG3. n = 6. **(D)** 2pSAM illustration of pneumococcal tracking in the spleens of WT and *Cd19*^-/-^ mice post infection with 5 × 10^6^ CFU of native *Spn*6A-GFP or that pre-opsonized with 5 µg T15 IgM (*Cd19*^-/-^ + T15, Movie S5). Scale bar, 50 μm. (C). Number of trapped pneumococci in the field during the first hour as monitored by 2pSAM in (C). n = 4 random FOVs. **(F-H)** Dose-dependent immune protection by anti-PC antibodies against pneumococcal infection. Bacteremia kinetics in the first 12 hr (F), at 24 and 48 hr (G), and survival (H) were assessed in WT mice post infection with 10^6^ CFU of *Spn*6A that untreated or pre-opsonized with low (1 µg) or high (50 µg) dose of T15 IgM. n = 6. Data were representative results (A, D, and E) or pooled (B, C, and F to H) from two independent experiments. Significance was compared by one-way (A) or two-way (B, C, and E to G) ANOVA and log-rank test (H). * *P* < 0.05, ** *P* < 0.01, *** *P* < 0.001, **** *P* < 0.0001.

To test whether RP macrophages depend on anti-PC antibodies to eliminate HV pneumococci, we performed a passive protection experiment in *Cd19^-/-^*mice with purified nAbs that were pre-absorbed with PC-coated resin. Pre-opsonization with PC-absorbed nIgM showed weakened activity in promoting *Spn*6A clearance in *Cd19^-/-^* mice as compared with the native nIgM, particularly at 3 and 6 hpi (**Fig. 5B**). To verify the specific role of anti-PC antibodies, we generated the IgM and IgG3 forms of anti-PC monoclonal antibodies of the T15 idiotype ^49^, which have been known to protect mice against pneumococcal disease ^50,51^. Opsonization with both T15 IgM and IgG3 potently rescued pneumococcal clearance in *Cd19*^-/-^ mice even at a low dose, e.g. 5 μg (**Fig. 5C**). We next monitored the *in vivo* behavior of pneumococci in the presence of T15 antibodies using 2pSAM. The T15 IgM was chosen because of its higher levels in antigen binding (**Fig. S5C**) and in promoting pneumococcal clearance (**Fig. 5C**). As expected, the long-term observation revealed sporadic *Spn*6A-GFP that trapped in the spleen of *Cd19*^-/-^ mice; this functional deficiency was fully recovered by pre-opsonization with T15 IgM (**Fig. 5D and 5E; Movie S5**). These functional investigation and real time illustration demonstrated that RP macrophages utilize anti-PC nAbs to clear the liver-resistant pneumococci. The C-reactive protein (CRP) has long been known as an important pentraxin that reacts with the PC moiety of pneumococcal C polysaccharide ^52^; however, the *Crp*^-/-^ mice were still able to clear the HV *Spn*6A in the first 12 hpi (**Fig. S5D**). This result indicated an essential role of nAbs in the splenic control of the liver-resistant pneumococci.

Since the importance of the anti-PC nAbs in clearing HV pneumococci, we next determined whether supplementation of anti-PC antibodies in normal mice could improve the bacterial clearance and rescue fatal infection. Pre-opsonization with monoclonal T15 IgM led to accelerated *Spn*6A clearance from the blood of WT mice in a dose-dependent manner. Whereas pretreated with a low dose (1 μg) of T15 IgM resulted in mildly enhanced bacterial clearance in the first 12 hpi, the blood bacteria were rapidly removed when using a high dose (50 μg) of T15 IgM as early as in 1 hpi (**Fig. 5F**). The blood bacteria were maintained at low levels in 2 days in the mice infected with extensively opsonized *Spn*6A, compared to the regrowth of bacteria at later stage without or with the low-dose antibody treatment (**Fig. 5G**). Accordingly, the high-dose antibody treatment led to complete elimination of *Spn*6A in half of the mice that were rescued from the lethal infection (**Fig. 5H**). These data suggested the anti-PC antibodies as a potential therapeutic choice to treat blood infection caused by HV pneumococci.

### Complement C3 is necessary for the splenic clearance of HV pneumococci

In the context of the abovementioned data, we investigated the potential involvement of the complement system in activating the anti-HV pneumococcal immunity of RP macrophages using *C3*^-/-^ mice ^53^, which lack the core complement protein C3. Similar to the RP macrophage- and antibody-deficient mice, the *C3*^-/-^ animals showed significant impairment in clearing *Spn*6A pneumococci from the circulation. The *C3*^-/-^ mice showed remarkably higher levels of bacteremia at all of the tested time points in the first 12 hpi with 10^6^ CFU of *Spn*6A, as compared with WT (**Fig. 6A**). Accordingly, the blood bacterial burden at 24 hpi in *C3*^-/-^ mice was 1,700-fold higher than the WT (**Fig. 6B**), leading to the 100% mortality in 36 hpi (**Fig. 6C**, left panel). Notably, all of the tested *C3*^-/-^ mice succumbed to the infection with 10^3^ CFU of *Spn*6A, an otherwise nonlethal dose (**Fig. 6C**, right panel). In addition, the cultured primary RP macrophages showed a significant level of binding to *Spn*6A bacteria that were pre-opsoinzed with normal serum, while the bacterial binding was significantly decreased when using *C3*^-/-^ serum for opsonization (**Fig. 6D**). These results strongly suggested that the complement-mediated innate immunity is involved in clearing the HV pneumococci in the early phase of septic infection.

**Figure 6.**
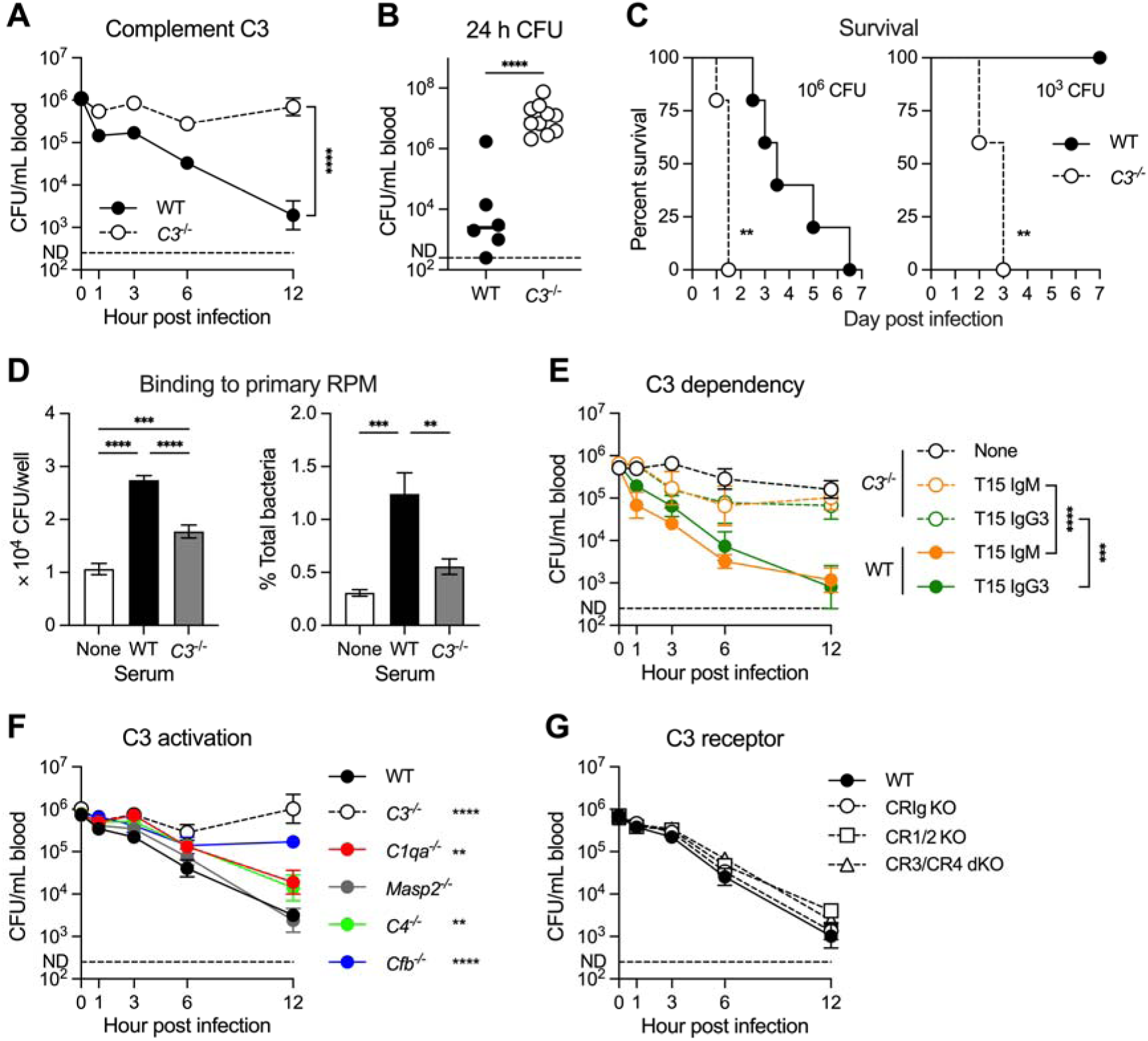
The requirement of complement C3 for the clearance of HV pneumococci in the spleen. **(A-C)** Pivotal role of complement C3 for the innate immune defense against HV pneumococci. Blood bacterial loads were counted in the first 12 hr (A) and at 24 hr (B) post infection with 10^6^ CFU of *Spn*6A in WT and *C3*^-/-^ mice. n = 6-12. Survival (C) of the WT and *C3*^-/-^ mice were monitored post infection with high dose (10^6^ CFU, left panel) or low dose (10^3^ CFU, right panel) of *Spn*6A. n = 5-6. **(D)** *In vitro* binding of HV pneumococci to primary RP macrophages. *Spn*6A was pre-opsonized by WT or *C3*^-/-^ serum and incubated with the primary RP macrophages for 30 min at MOI of 1. Bacterial binding levels were presented as the CFU numbers (left panel) and percentage (right panel) of cell-associated bacteria out of the total bacteria. n = 3. **(E)** Requirement of complement C3 for anti-PC antibody-mediated clearance of HV pneumococci. Blood bacterial loads were counted in the first 12 hr in WT and *C3*^-/-^ mice post infection with 10^6^ CFU of *Spn*6A that pre-opsonized by 5 µg of T15 IgM or IgG3. n = 6. **(F)** Role of complement activation pathways in the immune clearance of HV pneumococci. Blood bacterial loads were compared in the first 12 hr in each mouse line post infection with 10^6^ CFU of *Spn*6A. The *C1qa*^-/-^, *Masp2*^-/-^, and *Cfb*^-/-^ mice were specifically deficient in the classical, lectin, and alternative pathways; *C4*^-/-^ mice were inactive in both classical and lectin pathways. n = 6. (E). Dispensable role of well-known complement receptors in the immune clearance of HV pneumococci. Blood bacterial loads in complement receptor-deficient mice were assessed as in (E). Genetically knockout mice in CRIg (*Vsig4*^-/-^), CR 1/2 (*Cr1/2*^-/-^), and CR3 + CR4 (*Cd11b*^-/-^*Cd11c*^-/-^) were used. n = 6. Data were representative results (D) or pooled (A to C and E to G) from two independent experiments. Significance was compared by two-way (A, E, and F) or one-way ANOVA (D), student’s t (B), or log-rank (C) test. ** *P* < 0.01, *** *P* < 0.001, **** *P* < 0.0001.

To test the direct contribution of complement system on the anti-pneumococcal function of nAbs, we evaluated the bacterial clearance in *C3*^-/-^ mice post infection with T15 IgM- or IgG3-coated *Spn*6A. Although the anti-PC monoclonal antibodies promoted pneumococcal clearance from the bloodstream at 3 and 6 hpi, the bacteremia levels were still 10 to 20-fold higher in *C3*^-/-^ mice than in WT controls. Notably, the *C3*^-/-^ mice sustained a prominent bacterial burden in the blood during the first 12 hpi, despite the presence of T15 antibodies, compared with gradual clearance in WT mice (**Fig. 6E**). This *in vivo* assay demonstrated a pivotal role of the complement system in natural antibody-driven innate defense against HV pneumococci.

We next determined how natural antibody and the complement system work together to enhance the clearance of HV *S. pneumoniae* by RP macrophages. NAbs is known to activate the C3 protein via the classical activation pathway, with the help of complement proteins C1 and C4 ^43^. We thus tested the contribution of C1 and C4 to pneumococcal clearance using *C1qa*^-/-^ and *C4*^-/-^ mice ^54^. To our surprise, both the mouse lines showed marginal defect in eliminating *Spn*6A in the first 12 hr. Surprisingly, the *Cfb*^-/-^ mice lacking the alternative pathway activator protease factor B showed the most severe impairment in bacterial clearance (**Fig. 6F**), although there is no known functional linkage between antibody and factor B in C3 activation. C3 is known to promote bacterial phagocytosis by binding to complement receptors on phagocytes ^55^. The previous study has shown that RP macrophages express both complement receptor 3 (CR3) and 4 (CR4) ^32^. However, simultaneous knock out of CR3 and CR4 did not result in apparent impact on the *Spn*6A clearance during the first 12 hpi (**Fig. 6G**). Together, these data have revealed an essential role of C3 in mediating natural antibody-driven bacterial clearance by RP macrophages in the spleen, but the precise mechanism of the antibody-complement functional linkage awaits further investigation.

### The splenic anti-bacterial immunity operates across pneumococcal serotypes

Because PC is a conserved surface moiety in *S. pneumoniae*, we tested if anti-PC natural antibody-mediated splenic immunity operates across capsular serotypes by assessing the bacteremia kinetics of HV serotypes 2, 3, 4 and 8. All the four serotypes were gradually cleared from the bloodstream in the early phase of infection, although the clearance rates varied among the serotypes (**Fig. S6A**). However, the clearance immunity was completely lost by surgical removal of the spleen. As seen with *Spn*6A (**Fig. 1E**), asplenic mice showed striking increase in bacteremia level at various time points in the first 12 hr post infection with serotypes 2, 3, 4 and 8 (**Fig. 7A**). In contrast, LV serotypes 19F and 23F bacteria were rapidly eliminated in both the asplenic mice and sham surgery controls in the first 1 hr post inoculation and remained undetectable ever since (**Fig. S6B and S6C**). This information has revealed that the spleen is able to eliminate multiple liver-resistant pneumococcal serotypes.

**Figure 7.**
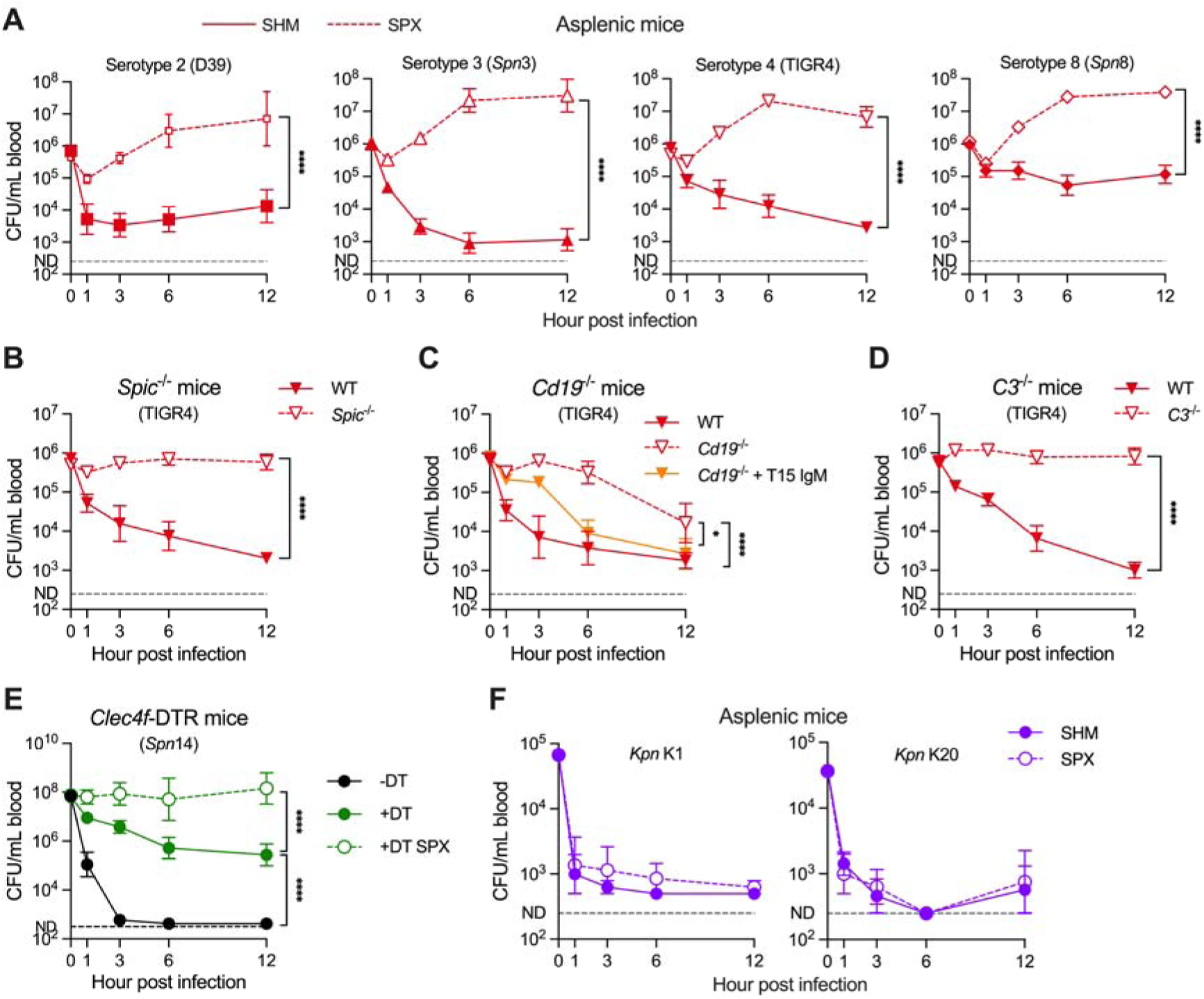
Serotype-independent broad operation of splenic RP macrophage-mediated immunity against HV pneumococci. **(A)** Importance of spleen for the blood clearance of multiple HV pneumococci. Blood bacterial loads in SPX mice were counted in the first 12 hr post infection with 10^6^ CFU of serotype 2 (D39), serotype 4 (TIGR4), and isogenic serotype 3 and 8 strains (*Spn*3 and *Spn*8), as compared with the SHM controls. n = 6. **(B)** Essential role of RP macrophages for the blood clearance of HV TIGR4. Blood bacterial loads in *Spic*^-/-^ mice were counted in the first 12 hr post infection with 10^6^ CFU of TIGR4 and compared with the WT controls. n = 6. **(C)** Promotion of anti-PC antibodies to the blood clearance of TIGR4. Blood bacterial loads in *Cd19*^-/-^ mice were counted in the first 12 hr post infection with 10^6^ CFU of the bacteria that untreated or pre-opsonized by 5 µg of T15 IgM. n = 6. **(D)** Pivotal role of complement C3 for the blood clearance of TIGR4. Blood bacterial loads in *C3*^-/-^ mice were counted in the first 12 hr post infection as in (B). n = 6. **(E)** Role of spleen in the clearance of LV pneumococci in liver immunocompromised mice. Blood bacterial loads in KC-specific depletion *Clec4f*-DTR (+DT), control (-DT), and KC-depleted SPX (+DT SPX) mice were compared post i.v. infection with 10^8^ CFU of the LV *Spn*14. n = 6. **(F)** Dispensable role of spleen in the clearance of HV *K. pneumoniae*. Blood bacterial loads in SPX mice were counted in the first 12 hr post infection with 10^5^ CFU of serotype K1 (left panel) or 5 × 10^4^ CFU of K20 (right panel) strains, and compared with the SHM controls. n = 6. All data were pooled from two independent experiments. Significance was compared by two-way ANOVA test. * *P* < 0.05, *** *P* < 0.001, **** *P* < 0.0001.

We further verified the cellular and molecular mechanisms of the broad anti-bacterial immunity in the spleen using strain TIGR4 (serotype 4). Consistent with the data obtained with *Spn*6A (**Fig. 2B**), RP macrophage-deficient mice displayed persistent bacteremia in the first 12 hpi (**Fig. 7B**). Moreover, the antibody-deficient *Cd19*^-/-^ mice exhibited remarkable deficiency in clearing TIGR4, particularly at the early time points, which could be reversed by pre-opsonizing the bacteria with the T15 antibody (**Fig. 7C**). In a similar manner, inoculation of TIGR4 pneumococci in the mice lacking complement C3 led to uncontrolled bacterial growth in the bloodstream at the early stage (**Fig. 7D**). These findings were verified with serotype-2 strain D39 (**Fig. S6D-F**). These data allowed us to conclude that RP macrophages, in concert with anti-PC nAbs and complement C3, are pivotal for splenic removal of various liver-resistant pneumococci.

Patients with chronic liver diseases are at higher risk in developing bacterial sepsis ^56,57^. We thus tested if the spleen clears LV pneumococci in case the liver-based immunity is compromised using the LV serotype-14 pneumococci (*Spn*14). In line with the importance of liver-resident macrophages in eliminating LV pneumococci ^28^, KC-deficient mice (*Clec4f*-DTR mice treated with diphtheria toxin, +DT) showed remarkable deficiency in removing *Spn*14 in the first 12 hr post i.v. infection, as compared with the untreated mice (-DT) (**Fig. 7E**). Nevertheless, there was a gradual clearance of the blood *Spn*14 in KC-deficient mice in the first 12 hr, resulting in a 230-fold reduction in blood bacteria compared to the inoculum; but this slow clearance was completely lost after the spleen was surgically removed in DT-treated *Clec4f*-DTR mice (+DT SPX) (**Fig. 7E**). This result showed that the spleen is a vital backup firewall against encapsulated pneumococci that cannot be cleared by the liver.

We finally assessed if the spleen plays a role in the clearance of other encapsulated virulent bacteria by performing blood infection with HV serotype K1 and K20 strains of *K. pneumoniae* ^29^. Similar to the clearance pattern of HV pneumococci, normal mice showed substantial clearance of HV *K. pneumoniae* in the early phase of blood infection (**Fig. 7F**). However, surgical removal of the spleen or deficiency in RP macrophages did not result in significant impact on bacteremia kinetics (**Fig. 7F**). This data suggested that the splenic RP macrophage-dependent function is specific for the immunity to the PC-containing pneumococci but not other blood-borne bacteria.

## DISCUSSION

It has been well documented that humans lacking a functional spleen are highly susceptible to invasive infection by *S. pneumoniae* and other encapsulated bacteria ^58^; however, it remains largely unknown how the largest immune organ executes its innate immunity against these bacteria. In this study, we have uncovered the dominant role of RP macrophages in the splenic clearance of HV pneumococci by taking advantage of their ability to escape the liver surveillance ^28,29^. We’ve further demonstrated that this organ-specific innate immunity requires opsonization of nAbs targeting the capsule-covered antigens of *S. pneumoniae* and activation of the complement system. While RP macrophages, nAbs and the complement system are known to contribute to host innate immunity against invasive bacterial infections, this study represents the first mechanistic map of the anti-bacterial machinery in the spleen.

### The spleen is the backup immune organ for the hepatic anti-bacterial “firewall”

As represented by the “reticuloendothelial system” ^23,24^, there are a plenty of more recent evidence that both the spleen and liver are important in host defense against bacterial infections ^11,25,59–63^. However, it remains largely undefined how the two organs divide the labor in the clearance of invading bacteria. Our recent studies have defined that the liver KCs capture and kill unencapsulated and many LV encapsulated bacteria with astonishing efficiency in mice once they enter the blood circulation; the spleen is completely dispensable for the clearance of these bacteria ^28,29^. This work has shown that the spleen clears encapsulated bacteria that the liver fails to eliminate. On the one hand, the spleen eliminates the liver-resistant bacteria that are naturally refractory to the hepatic “firewall”. This is manifested by the survival and subsequent growth of the liver-resistant HV serotypes in the blood circulation in the absence of the spleen. On the other hand, the spleen is able to control the blood-borne LV encapsulated bacteria when the hepatic filtration system is compromised. The spleen became essential for clearing the otherwise liver-susceptible LV pneumococci in KC-depleted mice. In such scenarios, the spleen serves as a backup, albeit less efficient, immune organ of the hepatic anti-bacterial “firewall”.

In summary, the liver appears to serve as the default organ for removing the “conventional” microorganisms in the bloodstream (e.g. microbiota and LV encapsulated bacteria), while the spleen is reserved for dealing with the “emergency” conditions due to invasive infections of the liver-resistant pathogens or the impairment of the hepatic surveillance system. The layback role of the spleen, relative to the liver, in bacterial clearance is consistent with the well-known role of the spleen in sampling microbial antigens in the circulation for B cell activation and antibody production ^2^, which would provide more swift responses to potential future “emergency”. Technically, the recent availability of the liver resistant pneumococcal serotypes has enabled us to distinguish the unique contributions of the spleen and liver to bacterial clearance.

### RP macrophage is the major immune cell against the liver-resistant pneumococci in the spleen

Despite the importance of the spleen in host defense against microbial infections, the splenic immune cells that execute bacterial clearance are still sketchy and controversial. MZ macrophages bind pneumococcal capsules via the C-type lectin receptor SIGN-R1 ^14–17^, which has been suggested to be essential for the clearance of pneumococci in mice ^16,17^. In contrast, a more recent study has suggested RP macrophages, instead of MZ macrophages, as the leading phagocyte for phagocytic clearance of virulent *S. pneumoniae* in the spleen ^13^. *Leishmania donovani*-induced expansion of RP macrophages made mice more resistant to serotype-3 HV pneumococci although the parasitic infections also resulted in the simultaneous loss of MZ macrophages; in addition, neutrophils and dendritic cells are dispensable ^13^. A subsequent study has confirmed the role of RP macrophages in capture serotype-2 pneumococci in the spleen, but also showed that neutrophils are responsible for killing RP macrophage-trapped bacteria in the spleen ^11^.

This work has provided unequivocal evidence that RP macrophage is the major immune cell in the spleen that captures and kills the liver-resistant virulent pneumococci in the early phase of septic infection. Our conclusion is supported by the complete loss of pneumococcal clearance in RP macrophage-deficient mice. This phenotype was identified by HDL-mediated RP macrophage depletion, and later confirmed by SpiC-dependent genetic deficiency in RP macrophage development. In sharp contrast, parallel experiments targeting MZ and MP macrophages did not yield obvious impact on the splenic clearance of the liver-resistant bacteria. Moreover, we have found that neutrophils play a marginal role in the splenic clearance of virulent pneumococci in the early phase of pneumococcal infection. Splenic neutrophils dramatically increased at 3-6 hr post infection, but depleting neutrophils with 1A8 antibody did not yield significant impact on the eradication of blood-borne pneumococci. The *in vivo* imaging data also showed a dominant role of RP macrophages in capturing and killing pneumococci in the RP compartment of the spleen. It should be noted that neutrophils and inflammatory monocytes appear to confer a lower level of anti-pneumococcal immunity at 12 hr post infection, since Gr1 antibody-treated mice showed significant impairment in pneumococcal clearance from the bloodstream. In full agreement with our finding, Gerlini *et al.* have shown that splenic macrophages are much more important than neutrophils and inflammatory monocytes in the clearance of pneumococci from the blood circulation ^30^.

### The tissue resident macrophages shape the spleen- and liver-specific immune functions

In the context of our previous studies on the unique role of KCs in carrying out the liver-specific clearance of blood bacteria ^28,29^, defining the function of RP macrophage in the spleen-specific immunity against the liver-resistant pneumococci in this study has enabled us to understand how the spleen and liver carry the organ-specific anti-bacterial immunity. The RP macrophages and KCs share several anatomic and functional features ^64,65^. In contrast to the intrinsic mobility of certain tissue-resident macrophages (e.g. alveolar macrophage and peritoneal macrophage), both RP macrophages and KCs are immobilized to their niches and directly exposed to blood flow. RP macrophages and KCs have been well documented as highly phagocytic cells in taking up old/injured erythrocytes (for RP macrophages) and microbes (for KCs). RP macrophages and KCs each represent the most abundant macrophage population in the spleen and liver, respectively. Finally, both the macrophage types are characterized by their phagocytic capacities.

However, there are numerous fundamental differences between RP macrophages and KCs, which, to greater or less extents, contribute to the organ-specific immunity. KCs represent approximately 90% of all resident macrophages in the body, and are positioned at the gateway of the hepatic portal vein where blood from the gastrointestinal systems gather and pass through the liver sinusoids ^66^. The liver macrophages are also famous for its swift capture of circulating bacteria and other particles ^11,25,59–63,67–70^. A recent study has shown that KCs positioned near the entry of the portal vein are even more phagocytic than those located at the distal part of the hepatic vasculatures, so-called “commensal-driven immune zonation” ^61^. The anatomic position, exceptional abundance and enormous phagocytic capacity of KCs suit well with the default immune task of removing potentially abundant and harmful microbes and other particles from the blood circulation. Although RP macrophages are far lower in cell number than KCs, they outnumber the combination of all the other macrophage populations in the spleen of humans and mice ^64^. Moreover, RP macrophages are highly phagocytic as manifested by the effective uptake of aged/damaged erythrocytes or erythrophagocytosis ^71^.

In the light of the existing literature, resident macrophage-specific expression of bacterial pattern-recognition receptors is likely to be a major factor in defining the RP macrophage- and KC-specific activities in clearing blood-borne bacteria. In sharp contrast to the well-documented phagocytic activities of RP macrophage and KCs against bacteria, the receptors that recognize microbial molecules for macrophage phagocytosis are far less understood. However, our recent studies have shown that KCs capture the LV encapsulated bacteria by receptor-mediated recognition of capsular polysaccharides, as demonstrated by the requirement of asialoglycoprotein receptor (ASGR) for KC capture of serotype-7F and -14 pneumococci in the liver sinusoids ^28,29^. While ASGR represents the only known receptor for bacterial capsules on KC receptor, the liver macrophage also uniquely express the complement receptor CRIg for bacterial phagocytosis, which is virtually not expressed by any other immune cells beyond the liver ^60,63,67,72,73^. While there are no existing bacterium-specific receptors on RP macrophages, it is likely that these macrophages also require such receptors for capturing flowing bacteria in the bloodstream. The CD91 and CD163 scavenging receptors have been identified for the uptake of hemopexin-heme complexes and haemoglobin by RP macrophages, respectively ^74,75^.

### Anti-PC nAbs are essential for the splenic immunity against the liver-resistant *S. pneumoniae*

Since the 1980s, the protective effects of naturally occurred antibodies targeting non-capsular antigens in pneumococcal infection have been documented ^38,48,50^. NAbs of the T15 idiotype, which recognize the phosphocholine (PC) moiety in pneumococcal cell wall polysaccharides, have been proved to protect mice from lethal infection by multiple serotypes of pneumococci, with empirical evidence based on the phenotypic observation on blood bacterial clearance and post-infection survival rate ^51,76^. In this work, the antibody-null *µMT* mice and B1 cell-deficient *Cd19*^-/-^ mice revealed functional impairment in the early control of HV pneumococci, which could be dose-dependently reversed by purified nAbs from murine serum or recombinant monoclonal T15 antibodies. The essentiality of anti-PC nAbs was further confirmed by the *in situ* 2pSAM illustration of nIgM- and T15 IgM-recovered pneumococcal entrapment in the RP of *Cd19*^-/-^ mice. Our study has uncovered the cellular basis underlying the protection conferred by anti-PC nAbs, by which RP macrophages capture the circulating HV pneumococci in the spleen, thus maintaining a fundamental level of anti-pneumococcal immunity in the native host.

Our recent work has highlighted that vaccine-elicited anti-capsule antibodies strongly augment the immune clearance of HV pneumococci by liver KCs and sinusoidal endothelial cells, thus providing robust protection ^44^. Moreover, we found that the splenic defense can be bolstered by additional anti-PC antibodies, which could convert the slow bacterial clearance to a rapid elimination and partially protect the host from lethal infection, underscoring the viability of pneumococcal PC as vaccine candidates. Despite the success of current capsule-based pneumococcal vaccines ^77^, they do face the limitations in serotype coverage and the challenges of serotype replacement ^78^. Consequently, antibodies targeting the conserved non-capsular antigens offer a compelling strategy for broad-spectrum vaccines that are effective against all serotypes of *S. pneumoniae*. To this end, various pneumococcal surface proteins have been widely explored as vaccine antigens, with several protein-based vaccines being employed in clinical trials ^79,80^. However, there remains inter-strain variability in some pneumococcal surface proteins ^79,81^, which has been reported to promote immune evasion to variant-specific antibodies^82^. The strictly immutable nature of the pneumococcal cell wall PC makes it a promising antigen for future vaccine development. Nonetheless, ongoing research efforts are imperative to optimize the efficacy of immune clearance mediated by anti-PC antibodies and to explore their full clinical potential.

### Complement system is pivotal for the anti-bacterial immunity in the spleen

The complement system represents an important element of the innate immunity to bacterial infections, including *S. pneumoniae* ^83^. It is not surprising that complement systems played a role in the RP macrophage-dependent pneumococcal clearance. Indeed, our findings demonstrate that complement C3 is indispensable for the efficient elimination of HV pneumococci during the early stage of septic infection, acting synergistically with RP macrophages and nAbs to mediate bacterial clearance. Moreover, the monoclonal T15 IgM- or IgG3-opsonized pneumococci cannot be efficiently eliminated in *C3*^-/-^ mice as compared with the rapid clearance in WT controls. These results indicate the complement system as a pivotal downstream effector mechanism of natural antibody-driven innate defense in the spleen. Unexpectedly, the splenic immunity seems to be less dependent on the classical pathway initiated by natural antibody, because the *C1qa*^-/-^ mice held the capacity to eliminate HV pneumococci as the WT controls. Instead, we noticed a severe impairment in bacterial clearance in alternative pathway-deficient *Cfb*^-/-^ mice, which implies that the efficient capture of HV pneumococci on RP macrophages demands massive C3 deposition via the amplification through alternative pathway ^54^. Although we have established that C3 is required for the natural antibody-mediated pneumococcal clearance, our *in vivo* assessment showed dispensable roles of the known complement receptors, including CR1/2, CR3, CR4, and CRIg. How C3-coated pneumococci are destroyed calls for further elucidations.

### Splenic immunity is specific against *S. pneumoniae* and likely other PC-containing bacteria

In clinical, asplenia and hyposplenism increase the susceptibility to overwhelming post-splenectomy infections (OPSI), especially those caused by three encapsulated bacteria, *S. pneumoniae*, *N. meningitidis*, and *H. influenzae*, all of which contain PC in surface structures ^5,58^. Notably, epidemiological surveys show that up to 90% of OPSI are caused by *S. pneumoniae* ^5^. The current study explains the specialized role of the spleen in defending against pneumococci and likely other PC-containing encapsulated bacteria. Our data showed that mice lacking a spleen or deficient in RP macrophage development and natural antibody production dramatically lost the innate clearance of HV pneumococci, while removal of the spleen in mice retained the capacity to eliminate HV *K. pneumoniae*, a non-PC encapsulated bacterium. This finding suggests that the splenic RP macrophages constitute a critical immune bottleneck of invasive pneumococcal infection, while alternative effector mechanisms contribute to clear other bacterial species. For instance, the complement-dependent formation of MAC on the outer membrane sensitizes the Gram-negative bacteria to a range of antimicrobial effectors beyond those afforded by splenic macrophages ^84,85^. In contrast, the RP macrophage-natural antibody-complement axis makes the spleen as the sole effective immune machinery against the liver-resistant *S. pneumoniae* in the host without prior immunization. In that light, the exclusive importance of spleen immunity against pneumococcal bacteremia provides a mechanistic explanation of the hyper-susceptibility of asplenia human individuals to *S. pneumoniae* over other encapsulated bacteria.

### Limitation of this study

We have investigated the cellular and molecular basis of splenic immunity to HV pneumococci in murine sepsis models, which should be interpreted with caution in the context of human disease. The ubiquity of PC-reactive nAbs in human serum ^86^ and their cross protection in mouse models ^48^ strongly suggest the anti-PC antibodies as a conserved immune response to pneumococcal infection across mammalian species. Nonetheless, the translational relevance of our findings needs further exploration.

## ACKNOWLEDGEMENTS

We thank the Tsinghua research platforms for assistance in animal experimentation (Laboratory Animal Research Center), flow cytometry (Center for Biomedical Analysis), IVM imaging (Center for Cell Biology), and protein mass spectrometry (Center for Proteomics). This work was supported by the National Key R&D Program of China to H.A. and J.-R.Z. (2023YFC2308003 and 2023YFC2306300) and the National Natural Science Foundation of China to J.-R.Z. (82330071).

## AUTHOR CONTRIBUTIONS

Conceptualization, H.A. and J.-R.Z.; experimentation, H.A., Y.H., Z.Z., X.H., K.L., J.M. and X.T.; methodology, H.A., Y.H., Z.Z., H.Z. and J.W.; data analysis, H.A., Y.H., Z.Z., H.Z., J.W., Q.D. and J.-R.Z.; composition, H.A., Y.H., Q.D. and J.-R.Z.; funding, H.A. and J.-R.Z.

## SUPPLEMENTAL FIGURE LEGENDS

**Figure S1.**
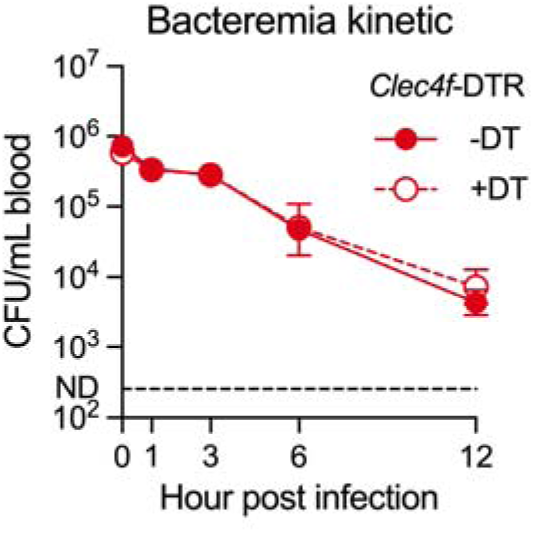
Dispensable role of liver KCs in clearing liver-resistant pneumococci. Bacteremia kinetics were monitored in KC-specific depletion *Clec4f*-DTR mice (+DT) and control mice (-DT) post i.v. infection with 10^6^ CFU of the *Spn*6A. n = 6. Data were pooled from two independent experiments and represented as mean ± SEM.

**Figure S2.**
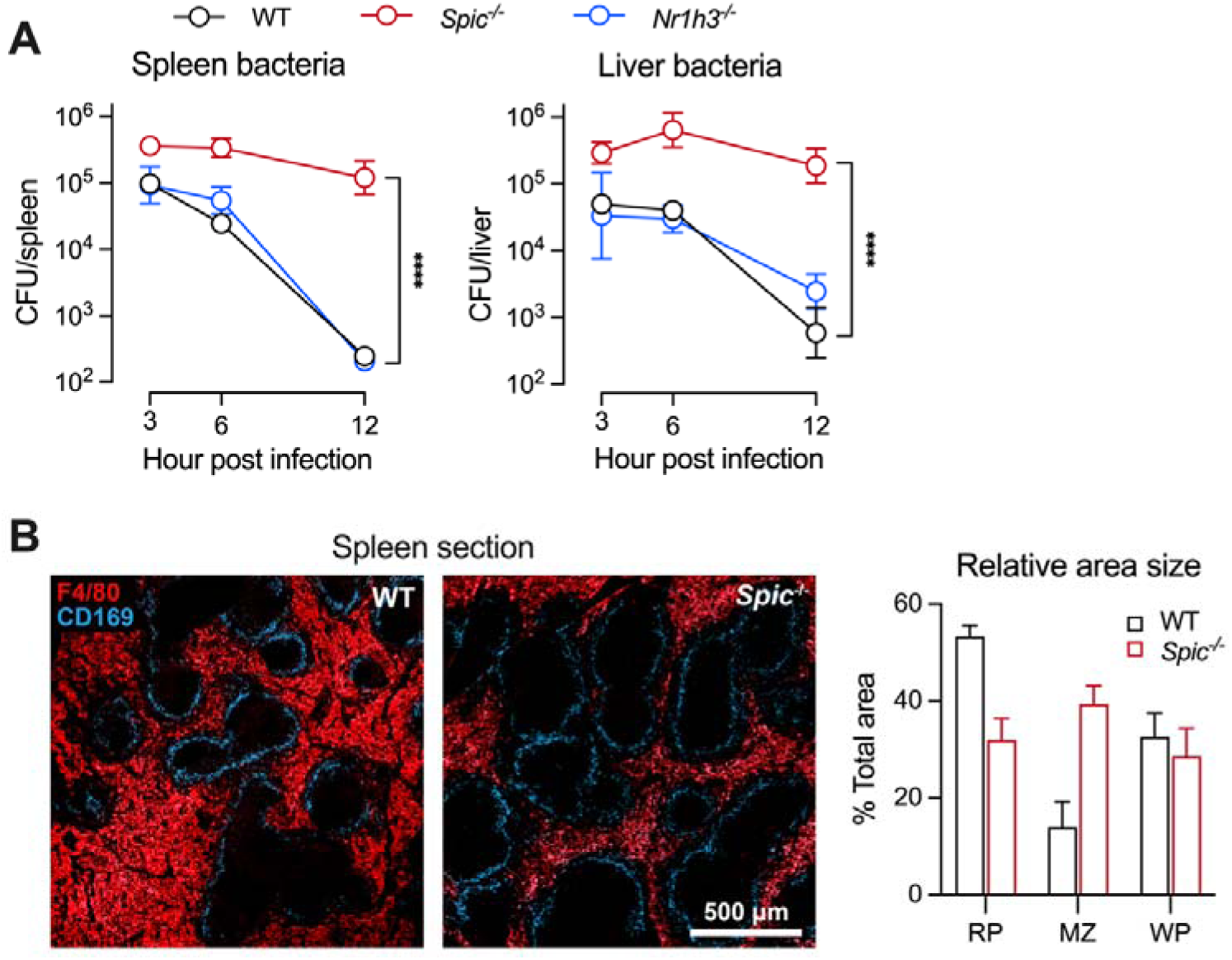
Essential role of RP macrophages in clearing HV pneumococci. **(A)** Essential role of RP macrophages in systemic control of HV pneumococci. Bacterial loads in the spleen and liver were counted in *Spic*^-/-^ or *Nr1h3*^-/-^ mice post infection with 10^6^ CFU of *Spn*6A. n = 6. **(B)** Representative fluorescence section to show the compartments of the spleen. RP and MZ were indicated by staining of AF647 F4/80 (Red) and AF594 CD169 (Blue), respectively. The WP (dark area) was delimited according to the MZ. Quantification of the area size of each compartment was calculated based on 5 sections and shown on the right. Data were pooled (A) or representative results (B) from two independent experiments and represented as mean ± SEM. Significance was compared by two-way ANOVA test. **** *P* < 0.0001.

**Figure S3.**
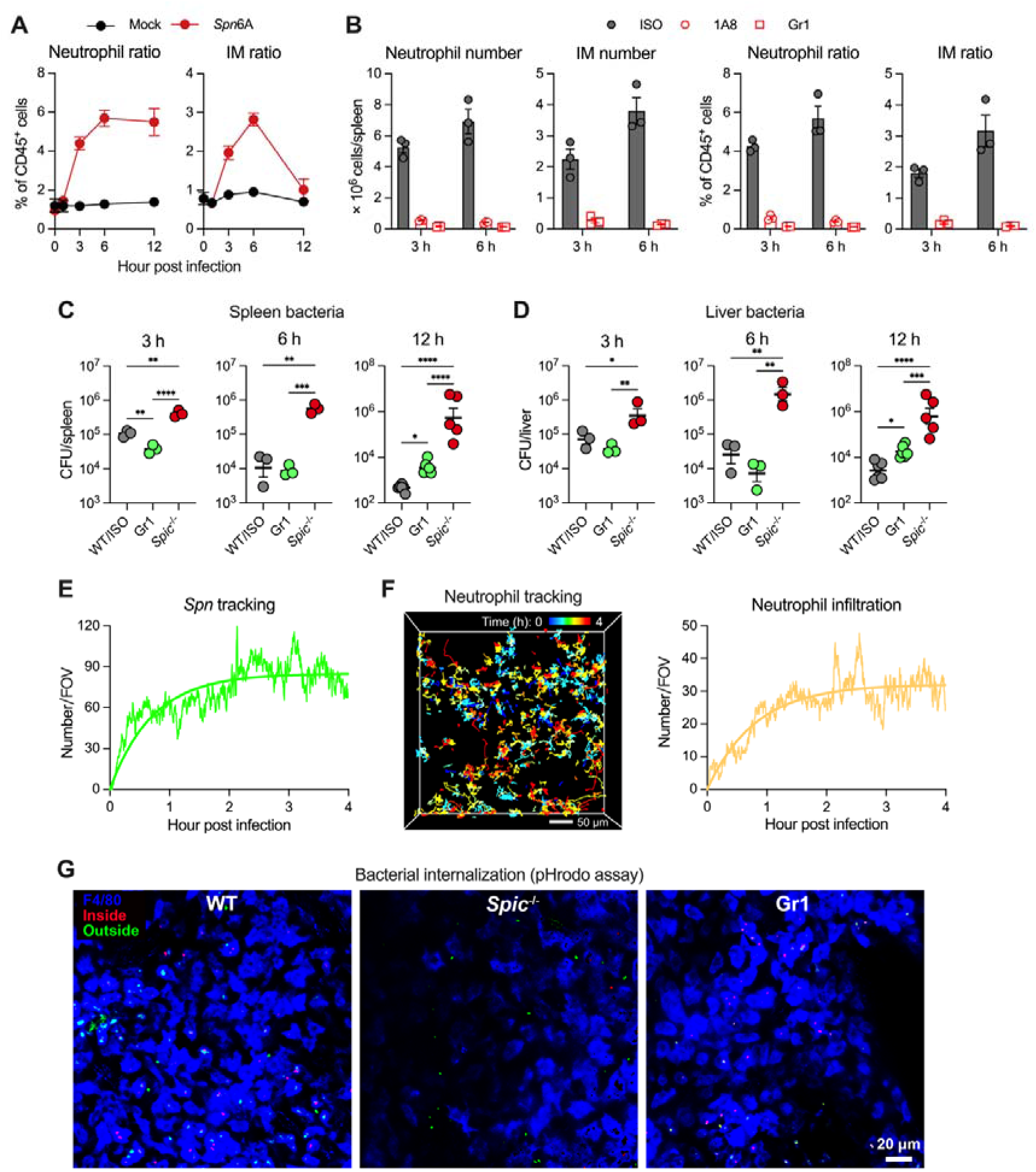
Neutrophil-independent elimination of HV pneumococci by RP macrophages. **(A)** Infiltration of neutrophils and IMs in the spleen during pneumococcal infection. The kinetics of ratio of neutrophils and IMs were assessed by flow cytometry post i.v. infection with 10^6^ CFU of HV *Spn*6A. Mock infected mice were i.v. injected with 100 μl of Ringer’s solution. n = 3. **(B)** Prevention of neutrophil and IM recruitment during the “eclipse phase” by antibody-mediated depletion. The mice were i.v. received with two doses of 500 ng 1A8 or Gr1 antibodies at 24 hr and 5 min prior to i.v. infection with 10^6^ CFU of HV *Spn*6A. The absolute numbers and ratio of neutrophils and IMs in the spleen were then assessed by flow cytometry. The same dose of isotype control (ISO) antibodies was included as controls. n = 3. **(C and D)** Impact of neutrophil and IM depletion on the systemic clearance of HV pneumococci. Bacterial loads in the spleen and liver were counted in control (WT/ISO), Gr1-treated, and *Spic*^-/-^ mice post infection with 10^6^ CFU of *Spn*6A. n = 6. **(E)** Curve of tracked pneumococcal cells during the first 4 hr post infection with 2 × 10^7^ CFU of GFP-expressing *Spn*6A (Movie S1). **(F)** Overlay of tracked traces (left panel) and curve of the quantity (right panel) of the infiltrated neutrophils during the first 4 hr post infection as in (A). Scale bar, 50 μm. **(G)** Representative 2pSAM images to illustrate the internalization of *Spn*6A in the spleen of WT, *Spic*^-/-^, and Gr1-treated mice. Uptake of *Spn*6A was indicated by the activation of pHrodo Red dye (Inside), whereas outside bacteria remained in green. Images were obtained at 0.5-1 hpi. Scale bar, 20 μm. Data were representative results (A, B, and E-G) or pooled (C and D) from two independent experiments and represented as mean ± SEM. Significance was compared by one-way ANOVA test (C and D). * *P* < 0.05, ** *P* < 0.01, *** *P* < 0.001, **** *P* < 0.0001.

**Figure S4.**
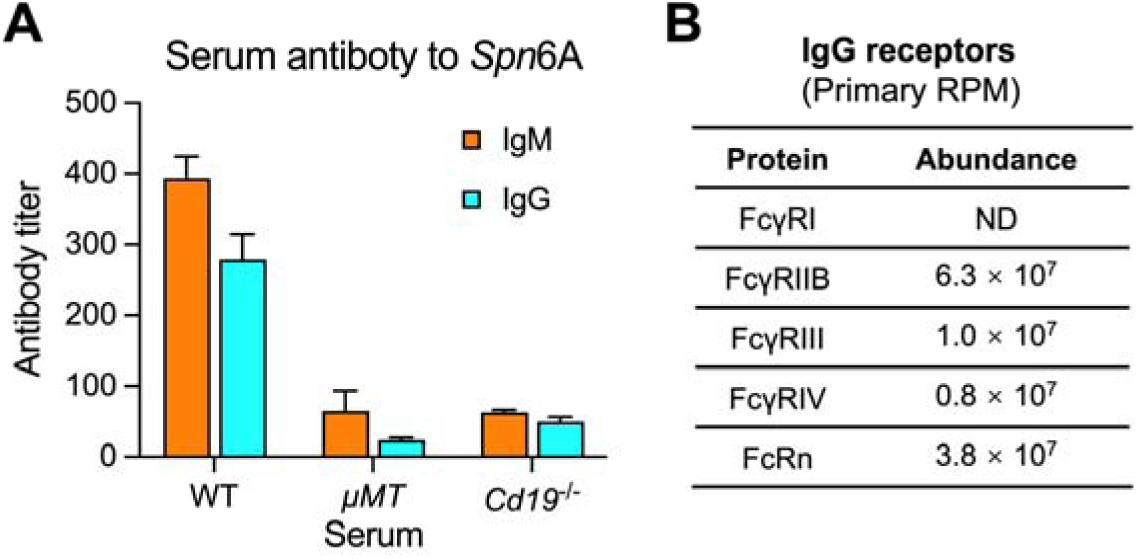
The essential role of natural antibodies in splenic clearance of HV pneumococci. **(A)** Detection of anti-pneumococcal natural antibodies in murine serum. Titers of IgM and IgG in serum isolated from WT, *µMT*, and *Cd19*^-/-^ mice were measured by ELISA using *Spn*6A whole cells as antigen. n = 3. **(B)** Relative abundance of known IgG receptors in murine RP macrophages. Data were the average amount of three replicates. ND, not detectable. Data in (A) were representative results from two independent experiments and represented as mean ± SEM.

**Figure S5.**
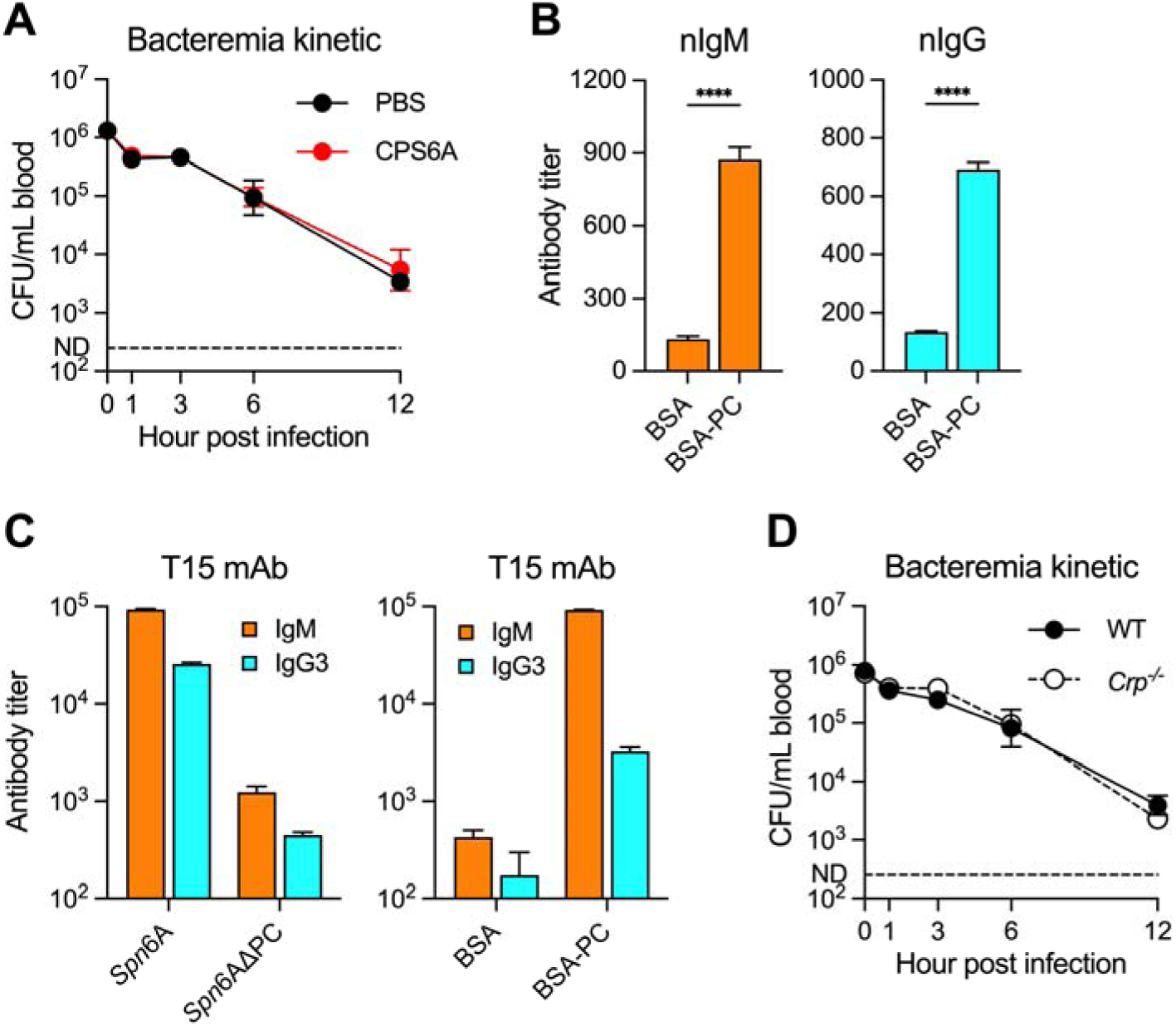
Pneumococcal cell wall PC as a target antigen of natural antibodies. **(A)** Irrelevance of capsular polysaccharide in HV pneumococcal clearance. Bacteremia kinetics were monitored in WT mice pre-administrated by PBS or 400 µg serotype 6A capsule (CPS6A) at 5 min before infection with 10^6^ CFU *Spn*6A. n = 6. **(B)** Detection of anti-PC antibodies in purified natural antibodies. Purified nIgM and nIgG were diluted at 1 mg/ml in PBS, and titers of IgM and IgG were measured by ELISA using BSA-conjugated PC as antigen. n = 3. **(C)** Verification of the specificity of the recombinant T15 monoclonal anti-PC antibodies. The T15 IgM and IgG3 were diluted at 1 mg/ml in PBS, and titers were measured by ELISA using *Spn*6A whole cells, PC-free *Spn*6A, or BSA-conjugated PC as antigens. n = 3. **(D)** Dispensable role of CRP in HV pneumococcal clearance. Bacteremia kinetics were monitored in WT and *Crp*^-/-^ mice post infection with 10^6^ CFU *Spn*6A. n = 6. Data were representative (B and C) or pooled (A and D) from two independent experiments and represented as mean ± SEM. Significance was compared by student’s *t* test (D). **** *P* < 0.0001.

**Figure S6.**
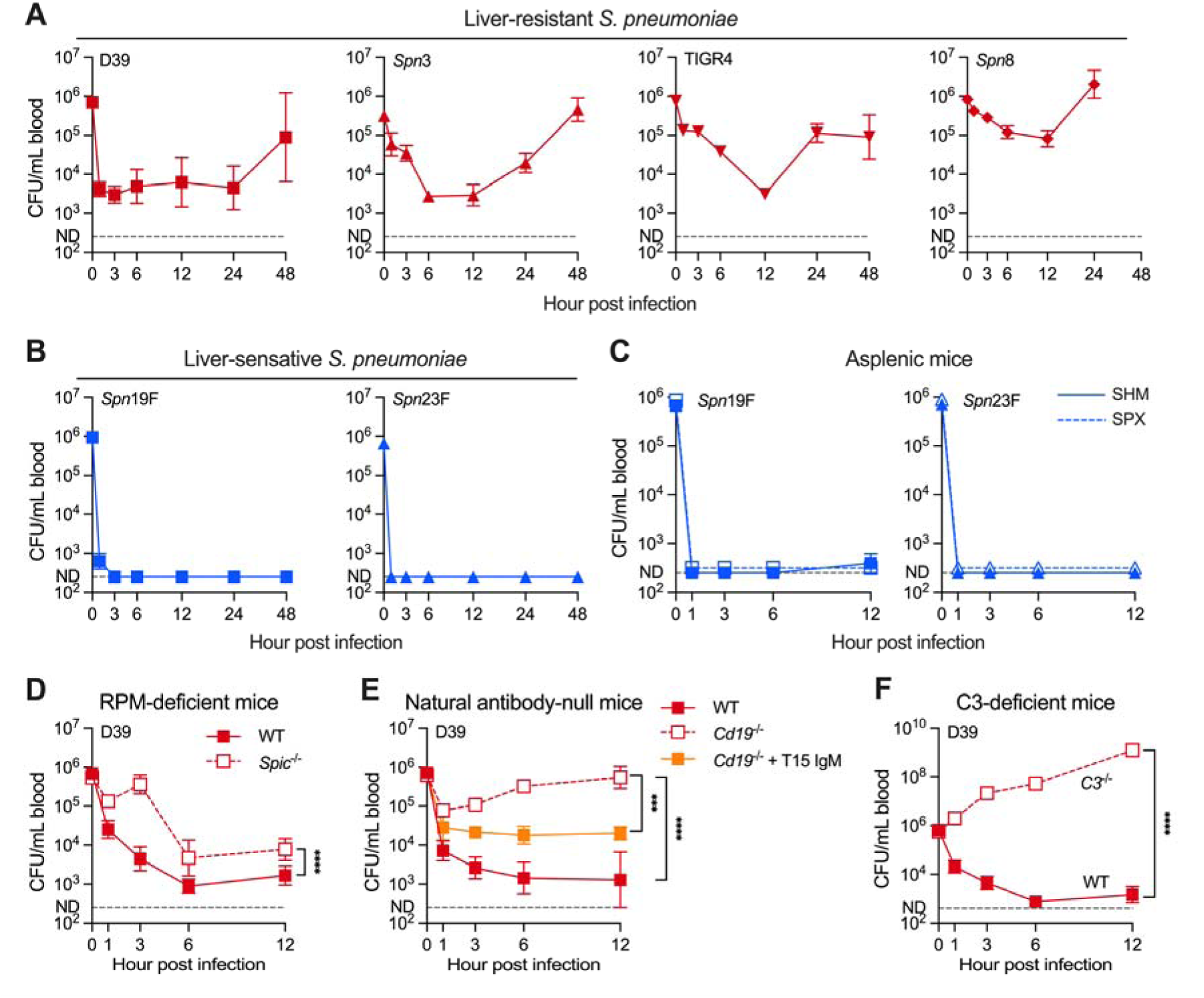
Serotype-independent splenic immunity against HV pneumococci. **(A and B)** Bacteremia kinetics in WT mice post i.v. infection with HV (A) and LV (B) pneumococci. Blood bacterial loads were counted post infection with 10^6^ CFU of HV serotype 2 (D39), serotype 4 (TIGR4), isogenic serotype 3 and 8 strains (*Spn*3 and *Spn*8), and LV serotype 19F and 23F strains (*Spn*19F and *Spn*23F). n = 6. **(C)** Dispensable role of spleen for the blood clearance of LV pneumococci. Blood bacterial loads in SPX mice were counted in the first 12 hr post infection as in (B), and compared with the SHM controls. n = 6. **(D)** Contribution of RP macrophages for the blood clearance of D39. Blood bacterial loads in *Spic*^-/-^ mice were counted in the first 12 hr post infection with 10^6^ CFU of D39 and compared with the WT controls. n = 6. **(E)** Promotion of anti-PC antibodies to the blood clearance of D39. Blood bacterial loads in *Cd19*^-/-^ mice were counted in the first 12 hr post infection with 10^6^ CFU of the bacteria that untreated or pre-opsonized by 5 µg of T15 IgM. n = 6. **(F)** Pivotal role of complement C3 for the blood clearance of D39. Blood bacterial loads in *C3*^-/-^ mice were counted in the first 12 hr post infection as in (D). n = 6. All data were pooled from two independent experiments and represented as mean ± SEM. Significance was compared by two-way ANOVA test (D-F). *** *P* < 0.001, **** *P* < 0.0001.

## MATERIALS AND METHODS

### Bacterial strains and cultivation

The *S. pneumoniae* and *K. pneumoniae* strains used in this study are described in **Table S1**. *S. pneumoniae* strains were cultured with Todd-Hewitt broth containing 0.5% yeast extract (THY) at 37L with 5% CO_2_, or on tryptic soy agar (TSA) plates supplemented with 5% defibrinated sheep blood as described ^87^. PC-free *S. pneumoniae* strains were obtained by cultivation of the bacteria in chemically defined medium (CDM) supplemented with 40 µg/ml ethanolamine instead of choline as described ^88^. *K. pneumoniae* strains were grown in Luria-Bertani (LB) broth or on LB agar plates as described ^29^.

### Mouse lines

The source and strain information of KO mice used in this work is provided in **Table S2**. The *Itgax*^-/-^ (CR4 KO) mice were inhouse generated by the CRISPR/Cas9 approach using single guide RNAs (gRNA15312) as listed in **Table S3**. The desirable offsprings were backcrossed with WT C57BL/6J mice for more than 7 generations before experimentation. The *Itgam*^-/-^*Itgax*^-/-^ (CR3/CR4 double KO) mouse was generated by CRISPR/Cas9 approach in *Itgam*^-/-^ background using sgRNA targeting *Itgax* locus. The resulting frameshifting mutations in *Itgam* and *Itgax* were confirmed by DNA sequencing. The *Vsig4*^-/-^*Itgam*^-/-^*Itgax*^-/-^ (CRIg/CR3/CR4 triple KO) mouse was obtained by crossing *Vsig4*^-/-^ and *Itgam*^-/-^*Itgax*^-/-^ mice.

### Septic infection

All animal experiments were performed in female C57BL/6 mice (6-8 weeks old) according to the animal protocols approved by the Institutional Animal Care and Use Committee in Tsinghua University (22-ZJR1). Septic infections were carried out by i.v. injection of desirable bacterial CFUs in 100 μl Ringer’s solution as described ^28^. The bacteremia kinetics were determined by retroorbital bleeding and CFU plating. Bacteria in the spleen and liver were measured by CFU plating of tissue homogenates and presented as CFU per organ. Mouse survival was recorded every day in a 7-day period or at a humane endpoint (body weight loss > 20%).

### Splenectomy

Asplenic mice were obtained by surgical removal of the spleen as described ^28^. In brief, mice were anesthetized with avertin (Sigma) at a dose of 400 mg/kg and meloxicam (Sigma) at a dose of 8 mg/kg before surgery. The peritoneum was opened on the left side to suture spleen pedicle before removing spleen and closing the peritoneum. Sham operated (SHM) mice underwent the same procedure without spleen removal. Post-operative animals were allowed to recover for at least 10 days before experiments.

### Immunofluorescence imaging of the spleen

Immunofluorescence staining of fixed spleen was carried out as described ^18^. Briefly, the spleens of mice infected with 10^7^ CFU of GFP-expressing *S. pneumoniae* were made into 10-µm frozen sections for confocal immunofluorescence imaging. RP and MZ macrophages were stained with F4/80-AF647 and CD169-AF594, respectively, at 3 µg/ml for 30 min after treatment with 200 µl blocking buffer (PBS supplemented with 1% BSA). Sections were mounted with an anti-fade mounting media after three washes in PBS. Images were acquired with Leica TCS SP8 confocal microscope using 10×/0.45 NA and 20×/0.80 NA HC PL APO objectives. The microscope was equipped with Acousto Optics without filters. Fluorescence signals were detected by photomultiplier tubes and hybrid photo detectors. Three laser excitation wavelengths (488, 585, and 635 nm) were employed by white light laser (1.5 mw, Laser kit WLL2, 470–670 nm). Representative images were acquired with 200× and 800× magnification at 1,024 × 1,024 pixels. The RP and MZ of the entire spleen section were photographed by concatenating multiple images at 100× magnification and the area was calculated by ImageJ (ImageJ 1.47v, National Institutes of Health, USA). Five to ten random FOVs were examined to calculate the number of captured bacteria in each area.

### Long-term 3D two-photon imaging of the spleen

Sub-cellular resolution imaging of the spleen was performed by 2pSAM essentially as described ^35^. Briefly, RP macrophages and neutrophils were stained with 2.5 µg of PE anti-F4/80 and PE-Cy5 anti-Ly6G, respectively, 30 min before i.v. infection with 5 × 10^6^ CFU of GFP-expressing *S. pneumoniae*. The spleen was surgically exposed and fixed by a vacuum apparatus under the lens. Mice were kept in anesthesia using 1.5% of isoflurane. An excitation light from a commercial femtosecond laser (Spectra-Physics InSight X3, Newport) was set at 1,000 nm for three-color imaging. The average laser power under the objective (25×/1.05 NA, water-immersion, Olympus, XLPLN25XWMP2) was about 30 mW. For detection, a 525-nm filter (MF525-39, Thorlabs) was used for bacteria imaging, a 610-nm filter (ET610/75m, Chroma) was used for macrophage imaging; a 670-nm filter (ET670/50m, Chroma) for the neutrophil imaging. Imaging data were collected at a 30 Hz sampling rate, and 512 pixels × 512 pixels × 13 angles scanning was adopted. Before 3D reconstruction, DeepCAD was used to perform denoising for each angle ^89^, and a customized denoising model was trained for each channel. The 3D reconstruction was performed as described ^35^.

Bacterial uptake by RP macrophages was visualized by 2pSAM as described ^28,90^. The splenic RP macrophages were stained by i.v. injection of 2.5 μg PE-Cy5 anti-F4/80 antibodies at 30 min before i.v. inoculation with 10^7^ CFU of pHrodo-labeled GFP-expressing *S.* pneumoniae. pHrodo labeling was conducted according to the manufacturer’s instructions except for using 0.001 mM pHrodo Red (Invitrogen). Images were acquired at 1 hr post infection.

Bacterial cell tracking was performed using a deep learning model that integrates the bacterial motility conditions and cellular feature similarity in 2pSAM images to comprehensively reconstruct cell movement trajectories as described ^91^. To reduce interference caused by extraneous bacterial and noise signals, bacteria appearing briefly (less than two frames) were excluded from the data.

### Immune cell depletion

MZ/MP macrophages and RP macrophages were selectively depleted by i.v. inoculation with low-dose (120 μg/mouse) and high-dose (1 mg/mouse) clodronate liposomes one day before infection, respectively, as described ^11^. Antibody-based depletion of neutrophils and inflammatory monocytes was accomplished as described in our previous study ^28^. KCs were removed by intraperitoneal injection of recombinant diphtheria toxin (DT) in *Clec4F*-DTR mice one day before infection ^28,92^.

### Flow cytometry

Flow cytometry was performed essentially as described ^28^. Total splenocytes were harvested by passing through a 70-µm cell strainer, and red blood cells were lysed by 1 ml RBC lysis solution. The splenocytes (10^6^) were blocked with 50 μl FACS buffer (PBS with 3% FBS) containing 1% anti-CD16/32 antibody for 10 min, and then stained with antibodies: APC-Cy7 anti-CD45 (1/200), BV605 anti-CD11b (1/500), FITC anti-F4/80 (1/200), PE anti-Ly6G (1/500), and PB anti-Ly6C (1/500) for 20 min. Cell viability was characterized by adding 5 μl of 7-AAD to the samples prior to analysis. Viable cells were gated as splenic RP macrophages (CD45^+^CD11b^+^F4/80^high^), neutrophils (CD45^+^CD11b^+^Ly6C^+^Ly6G^high^), inflammatory monocytes (CD45^+^CD11b^+^Ly6C^high^Ly6G^−^).

### Purification of nAbs

Natural IgM and IgG were purified from normal murine serum using Protein G and Protein L resin (GenScript, China) according to the manufacturer’s instructions. Briefly, a total volume of 10 ml serum was diluted with 10 ml PBS, and incubated with 1 ml Protein G resin at 4L for 2 hr. The resin was washed with 5 ml PBS for four times and bound IgG was eluted with 10 ml of 0.1 M glycine, pH 2.5. The flow-through serum from Protein G columns was further mixed with 1 ml Protein L resin to purify IgM in a similar manner. Purified IgM and IgG were concentrated by ultracentrifugation using 30 kDa centrifugal filters (Millipore, USA), sterilized by 0.22 μm centrifugal filter (Corning, USA), and quantified by the BCA Assay Kit (Beyotime, China).

### Recombinant monoclonal antibody production

Murine monoclonal antibodies (idiotype T15) against phosphocholine were generated as described with minor modifications ^93^. The coding sequences of variable region for heavy chain (VH, GenBank accession M16334.1) and light chain (VL, GenBank accession U29423.1) were synthesized according to published sequences ^94,95^. The coding sequences of constant region for heavy chains (CH_IgM_, CH_IgG3_, CH_IgG1_), light chain (CL), and J chain were PCR amplified using murine spleen cDNA as templates. The full-length spleen cDNA was amplified from the total RNA extracted from mouse spleen by TRIzol Reagent (Invitrogen) using Maxima H Minus First Strand cDNA Synthesis Kit (Thermo Scientific). The DNA fragments were further linked by fusion PCR and cloned into H and L vectors by enzymatic digestions and ligations. The relevant primers and resulting plasmids are listed in **Tables S3** and **S4**. The full-length IgG3 and IgM antibodies were produced by co-transfection of the H and L vectors into the HEK293 suspension culture cells (Expi293F) and purified with Protein G (IgG) and Protein L (IgM) resin, respectively.

### ELISA

Antibody titers of murine serum and purified IgG/IgM were quantified by enzyme-linked immunosorbent assay (ELISA) as described ^96^. Briefly, antigens in 100 μl PBS were coated on 96-well plates at indicated concentrations: pneumococcal cells, OD_600 nm_ 0.1; BSA and BSA-PC, 10 μg/ml; CPS, 10 μg/ml. BSA-PC was produced as reported ^97^. Briefly, 75 mg of cytidine 5’-diphosphocholine (CDPC, Sigma) was oxidized in 2.5 ml of 0.1 M sodium periodate for 20 min before 0.15 ml of 1 M ethylene glycol was added to stop the reaction. BSA (140 mg, Sigma) was dissolved in 5 ml of 0.1 M sodium bicarbonate and incubated with the activated CDPC for 1 h. After that, 5 ml of 0.5 M sodium borohydride was added, and the mixture was incubated overnight. The resulting solution was buffer-exchanged with PBS by ultracentrifugation. CPS was extracted as described ^28^. Immunoglobulin class and subtype were determined with HRP-conjugated anti-mouse IgM, IgG1, IgG2b, IgG2c, and IgG3 antibodies (EasyBio, Beijing, China).

### Construction of antibody receptor-expressing CHO cells

CHO cell lines expressing mouse antibody receptors were constructed as described ^44^. For expression of mouse *Fcmr* and *Fcamr*, the full-length complementary DNAs (cDNAs) of the target mouse genes were amplified from the total RNA isolated from the spleen using TRIzol Reagent (Invitrogen) using Maxima H Minus First Strand cDNA Synthesis Kit (Thermo Scientific) and cloned into pCDH vector with a His_6_ tag at the C terminus. The ligation mixtures were transformed into *E. coli* DH5α and selected on LB plates with 100 μg/ml ampicillin. Recombinant plasmids were confirmed by DNA sequencing and extracted using HiPure Plasmid EF Micro Kit (Magen) for subsequent lentiviral transduction. Recombinant plasmids were transfected into HEK293T cells using Lipofectamine 2000 (Invitrogen) together with lentiviral packaging vectors pMD2.G and psPAX2 (gifts from Didier Trono, Addgene). The lentiviral particles were harvested at 48 h and filtered through 0.45 μm syringe filter unit (Millipore) to remove cell debris. The pCDH-lentivirus was used to infect CHO cells with 8 μg/ml Polybrene (Sigma). The transfectants were selected with 5 μg/ml puromycin for 7 days. The relevant primers and resulting plasmids are listed in **Tables S3** and **S4**.

### RP macrophage cultivation

Mouse primary RP macrophages we isolated and cultivated as described ^98^. In brief, the splenocytes were harvested by grinding the spleen and resuspended in 5 ml RPMI 1640 medium. Single cells were obtained by filtering through a 70-µm cell strainer and red blood cells were lysed with RBC lysis solution. Splenocytes from each spleen were resuspended in 10 ml of conditional medium (CM, RPMI 1640 supplemented with 20% L929 cell supernatant and 10% FBS) and cultured in a 10 mm plate at 37°C, 5% CO_2_ for 3 days. Nonadherent cells were removed, and the adhered RP macrophages were cultured in CM for another 4 days before use.

### *In vitro* bacterial binding

Bacterial binding to host cells were assessed essentially as described ^28^. CHO transfectants and primary RP macrophages were seeded in 96-well cell culture plates and grown to 90-100% confluence (∼ 5×10^4^ cells/well). At the time of binding experiments, growth media were replaced with basic F-12K (CHO cells) or RPMI 1640 (RP macrophages) without serum and antibiotics. For pre-opsonization, every 10^7^ CFU of pneumococcal cells were incubated with 20 μg of monoclonal T15 antibody or 50 μl mouse serum at 37°C for 30 min. Antibody or complement-coated bacteria were suspended in basic F-12K or RPMI 1640 at a density of 5×10^4^ CFU in 50 μl and added into 96-well plates with 50 μl per well, resulting a multiplicity of infection (MOI) of 1:1, followed by centrifugation at 500 g for 5 min to maximize the contact between bacteria and the cells. The mixtures were incubated for 30 min at 37°C with 5% CO_2_. The free bacteria were enumerated by CFU plating of the supernatants. The eukaryotic cells were thoroughly washed to remove free bacteria and lysed with 100 μl ice cold sterile H_2_O to enumerate cell-associated bacteria by CFU plating of the lysates. Bacterial binding was calculated by dividing the cell-associated CFU to the CFU of total bacteria.

### Quantification and statistical analysis

All experiments presented in this work were repeated at least twice at different times. The data are analyzed and presented as mean ± standard error of mean (SEM). Statistical analysis was performed using GraphPad Prism software (8.3.1). The levels of statistical significance are defined by *P* values of < 0.05 (*), < 0.01 (**), < 0.001 (***), and < 0.0001 (****). Flow cytometry and gene/protein sequence data were analyzed using FlowJo (10.4) and Lasergene (15.0.0) for Macintosh, respectively.

## SUPPLEMENTAL INFORMATION

**Table S1.** Bacterial strains.

**Table S2.** Mouse lines.

**Table S3.** Oligonucleotides.

**Table S4.** Recombinant plasmids.

**Movie S1.** Long-term illustration of *Spn*6A sequestration in the spleen of WT mouse by 2pSAM.

**Movie S2.** Comparison of *Spn*6A capture in the spleen of WT, *Spic*^-/-^, and Gr1-treated mice.

**Movie S3.** Illustration of neutrophil-dependent and independent elimination of RP macrophage-captured *Spn*6A.

**Movie S4.** Essential role of natural antibodies in *Spn*6A clearance in the spleen.

**Movie S5.** Promotion of *Spn*6A capture in the spleen by monoclonal anti-PC IgM.

