## Supplementary material for "Splenic red pulp macrophages eliminate the liver-resistant *Streptococcus pneumoniae* during bloodstream infection": Table S1

**Table S1. Bacterial strains used in this study**

| Strain ID | Species | Serotype | Description |
| --- | --- | --- | --- |
| D39 | *S. pneumoniae* | 2 | Wild type |
| TIGR4 |  | 4 | Wild type |
| TH870 |  | 6A | Wild type (*Spn*6A) |
| TH15921 |  | N/A | Marker-free capsule mutant of TH870 (*Spn*6AΔCPS) |
| TH15939 |  | 3 | Isogenic serotype 3 strain in TH870 background (*Spn*3) |
| TH15943 |  | 8 | Isogenic serotype 8 strain in TH870 background (*Spn*8) |
| TH15944 |  | 14 | Isogenic serotype 14 strain in TH870 background (*Spn*14) |
| TH14188 |  | 19F | Isogenic serotype 19F strain in TH870 background (*Spn*19F) |
| TH15945 |  | 23F | Isogenic serotype 23F strain in TH870 background (*Spn*23F) |
| TH12908 | *K. pneumoniae* | K1 | Wild type (*Kpn* K1) |
| TH13092 |  | K20 | Wild type (*Kpn* K20) |
