## Supplementary material for "Splenic red pulp macrophages eliminate the liver-resistant *Streptococcus pneumoniae* during bloodstream infection": Table S3

**Table S3. Oligonucleotides used in this study**

| ID | Sequence (5’-3’) |
| --- | --- |
| sgRNA |  |
| gRNA15312 | GGTGGTGGTTGGAGCACCAA |
| Primer |  |
| Pr19033 | GCTCTAGAGCCACCATGGACTTTTGGCTTTGGTTACTTTACTTC |
| Pr19034 | CCGCTCGAGTCAATGGTGATGGTGATGATGTTGGCATGAAGATCTGGGCCCTGGG |
| Pr19035 | GCTCTAGAGCCACCATGGACCAAGGTGCCCCAGCTAAGCCCAGT |
| Pr19036 | CCGCTCGAGTCAATGGTGATGGTGATGATGGGGTCTTGGGTCATTCTCCAGGACG |
| Pr19047 | CTGGAAATCAAGCGTACGCGTACGGATGCTGCACCAACTGTAT |
| Pr19048 | GCGGCCAAGCTTGGGAGCGGCCGCTCAACACTCATTCCTGTTG |
| Pr19134 | GAGAACCGGTGTACATTCTGAGGT |
| Pr19135 | GAGACTCGAGGCTGAGGAGACGGT |
| Pr19136 | GAGAACCGGTGTACATTCTGACATTG |
| Pr19137 | GAGACGTACGCCGTTTCAGCTCC |
| Pr19182 | GGAATTCATGAAGAACCATTTGCTTTTCTG |
| Pr19183 | GGATGATACATGACCATCCCATAGGGCCGGGATTCTCCTC |
| Pr19184 | GAGGAGAATCCCGGCCCTATGGGATGGTCATGTATCATCC |
| Pr19185 | GAGACTCGAGCAGTCAGTCCTTCCCAAATGT |
| Pr19186 | GAGAAAGCTTGGGAGCGGCCGCTCAATAGCAGGTGCCGCC |
| Pr19187 | GAGACTCGAGCCTCGAGCGCTACAACAAC |
| Pr19188 | GGGCCATGGCGGCCAAGCTT |
