## Supplementary material for "Splenic red pulp macrophages eliminate the liver-resistant *Streptococcus pneumoniae* during bloodstream infection": Table S4

| Plasmid ID | Backbone | Insertion segments | Primers for  segments | Template  DNA | Digestion |
| --- | --- | --- | --- | --- | --- |
| pTH16845 | Heavy chain vector for mouse IgM | J chain + VH T15 + CH_IgM_ | Pr19182/Pr19183 (J) | Mouse spleen cDNA | EcoRI/HindIII |
|  |  |  | Pr19184/Pr19135 (VH T15) | GenBank: M16334.1 |  |
|  |  |  | Pr19182/Pr19135(J-VH fusion) | J + VH |  |
|  |  |  | Pr19185/Pr19186(CH_IgM_) | Mouse spleen cDNA |  |
| pTH16838 | Heavy chain vector for mouse IgG3 | VH T15 + CH_IgG3_ | Pr19134/Pr19135 (VH T15) | GenBank: M16334.1 | AgeI/HindIII |
|  |  |  | Pr19187/Pr19188 (CH_IgG3_) | Mouse spleen cDNA |  |
|  |  |  | Pr19134/Pr19188 (VH T15- CH_IgG3_ fusion) | VH + CH_IgG3_ |  |
| pTH14793 | Light chain vector for mouse Ig | VL T15 + CL | Pr19136/Pr19137 (VL T15) | GenBank: U29423.1 | AgeI/NotI |
|  |  |  | Pr19047/Pr19048 (CL) | Mouse spleen cDNA |  |
|  |  |  | Pr19136/Pr19048 (VL T15-CL fusion) | VL + CL |  |
| pTH16693 | pCDH | Mouse Fcmr | Pr19033/Pr19034 | Mouse spleen cDNA | XbaI/XhoI |
| pTH16694 | pCDH | Mouse Fcamr | Pr19035/Pr19036 | Mouse spleen cDNA | XbaI/XhoI |

**Table S4. Construction of recombinant plasmids for antibody production**
